# Progressive neurodegeneration in human dorsal root ganglion from diabetes to painful neuropathy

**DOI:** 10.64898/2026.01.16.700028

**Authors:** Ishwarya Sankaranarayanan, Khadijah Mazhar, Allison M. Barry, Stephanie Shiers, Andi Wangzhou, Julia R. Sondermann, Feng Xian, Nikhil Inturi, Michael A. Wilde, Eric C. Meyers, Joseph B. Lesnak, Jayden A. O’Brien, Muhammad Saad Yousuf, Manuela Schmidt, Gregory Dussor, Erin Vines, Peter Horton, Anna Cervantes, Tariq Khan, Geoffrey Funk, Diana Tavares-Ferreira, PRECISION Human Pain Network, Patrick M. Dougherty, Michele Curatolo, Theodore J. Price

## Abstract

Diabetic painful neuropathy (DPN) is characterized by neuropathic pain accompanied by loss of sensory function. We hypothesized that neurodegeneration in the dorsal root ganglion (DRG) could underlie DPN progression. To address this question, we performed multi-omic evaluation on DRGs from otherwise healthy organ donors, donors with diabetes but no neuropathy, and donors with clinically diagnosed DPN. We discovered that the first stages of neurodegeneration begin early in diabetes before the onset of DPN, with Nageotte nodule formation accompanied by apoptotic gene expression and decreased proportion of specific populations of A-fibers in DPN with remodeling of non-neuronal cells. Our findings define DPN as a neurodegenerative disorder of the DRG, identify molecular markers of disease stage, and highlight the need for early intervention to prevent irreversible neurodegeneration.

## Main Text

DPN is the most common form of neuropathic pain, but its basic mechanisms in humans are still not completely defined and no specific, mechanism-based treatments for DPN are approved (*1–3*). A common feature of the disease is dieback of sensory axons from the skin of extremities (*1, 3, 4*) but the extent to which neurodegeneration occurs in the human DRG (hDRG) in DPN is still unknown. Recently we demonstrated that neuronal degeneration in the form of Nageotte nodule formation in the hDRG is associated with DPN (*5*), a finding that is consistent with bulk RNA sequencing experiments on DPN hDRGs (*6*). The purpose of this study was to understand the degree to which DPN is associated with neurodegeneration, molecularly characterize the progression of this degeneration from diabetes through advanced DPN, and identify specific neuronal types that are degenerating in DPN.

To achieve the above-stated goals, we recovered hDRGs from organ donors over a 5-year period carefully cataloging clinical characteristics of diabetes and DPN from medical records and family histories (**Supplementary Table 1**). We used a histological classification scheme coupled with a multi-omic approach that included single nucleus RNA sequencing (single-nuc RNAseq), Visium and Xenium spatial transcriptomics, and quantitative proteomics to comprehensively characterize how the progression of diabetes to DPN is reflected at the molecular level in the hDRG (**Fig. 1A**). We surveyed records from more than 400 donors and identified 25 donors with a history of diabetes or DPN, whose hDRGs were compared to 54 age- and sex-matched control donors, yielding a total of 79 hDRGs included in this study, of which 30 lumbar hDRGs were used for single-nucleus RNA sequencing. (**Fig. 1B**). Our findings provide strong evidence to support classification of DPN as a neurodegenerative disorder affecting the DRG that requires early intervention to prevent the advance of the disease.

**Figure 1.**
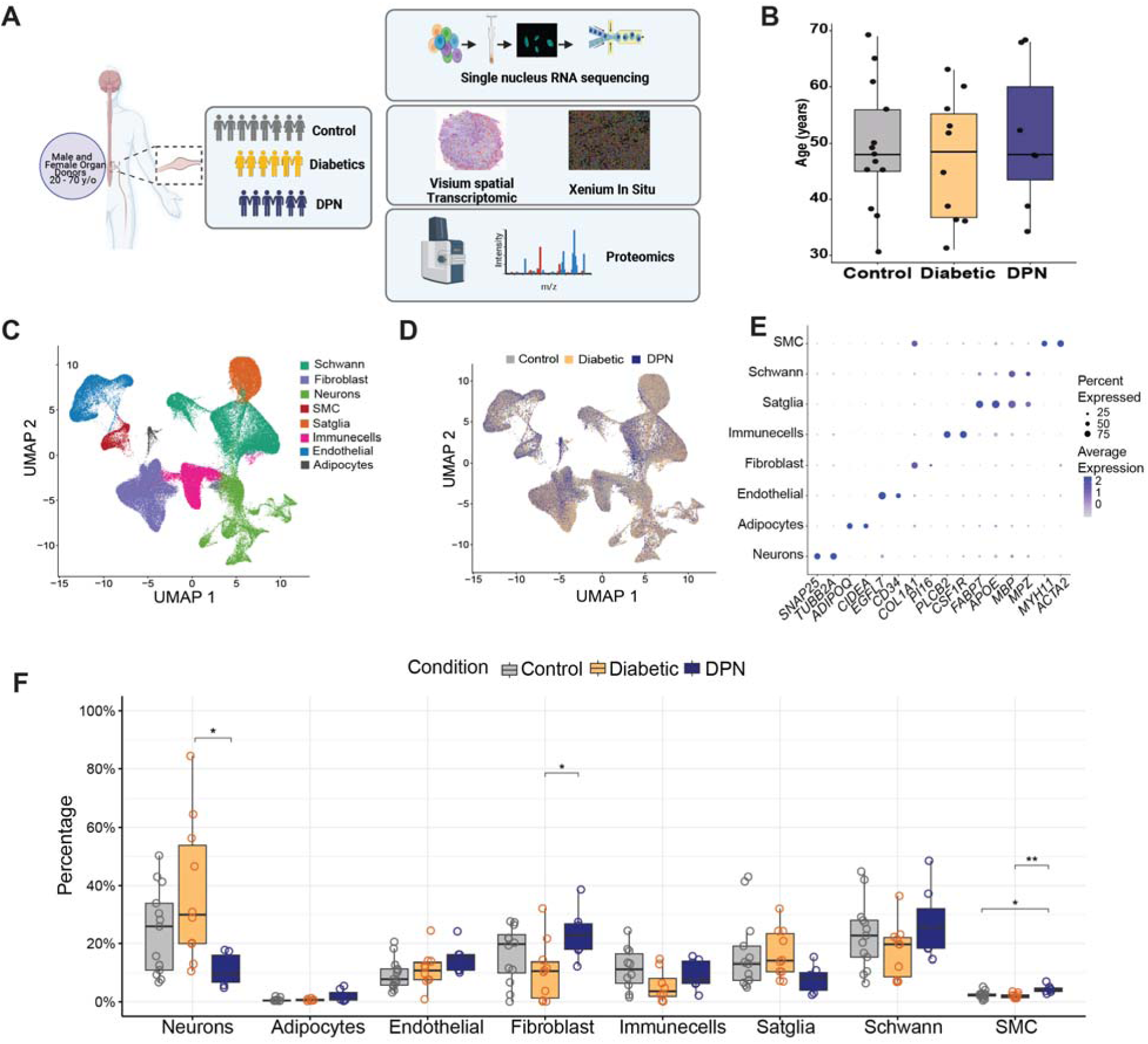
Single-nuc RNAseq profiling of hDRG in the progression of DPN reveals shifts in cell types loss. (**A**) Schematic of the multi-omics approaches applied to the human DRG cohort. (**B**) Bar plots showing the distribution of donor age and the number of donors included in the single-nucleus RNA-seq study. (**C**) UMAP projection of single-nucleus RNA-seq data showing the major cell types present in hDRG. (**D**) UMAP projection colored by donor condition (Control, Diabetic, and DPN). (**E**) Dot plots of representative canonical marker genes used to annotate major cell types. (**F**) Per-sample cell-type proportion analysis showing the percentage of cells per donor per condition, grouped by cell type. Each point represents one donor, with overlaid mean values per group. Statistical significance was assessed using Kruskal–Wallis tests followed by Dunn’s post hoc test with Benjamini–Hochberg correction for multiple comparisons; p < 0.05 was considered significant.

### Single cell characterization of diabetes to DPN progression in the hDRG

We conducted single-nuc RNAseq on 30 DRGs from 28 organ donors using the 10X Genomics FLEX RNA transcriptomics kits that allow for excellent recovery of hDRG neuronal nuclei. This resulted in sequencing of 172,740 total nuclei, including 13,121 neurons, across all 30 samples. We identified 8 cell types that were consistent with previous studies on hDRGs (**Fig 1C**). Nuclei from control, diabetic and DPN samples were represented in all the cell types across the UMAP projection (**Fig. 1D**). Canonical marker genes for each cell type were represented in appropriate clusters for each cell type (**Fig. 1E**). We examined the proportionality for each of the 8 cell types in control versus diabetic and DPN samples and found shifts occurring in the DPN cell types. We observed an increase in fibroblast and smooth muscle cell proportion and a decrease in neurons in DPN (**Fig. 1F and Supplementary Data 1**).

Given this finding of decreased neuronal proportion in DPN, and our previous description of increased Nageotte nodule formation in DPN DRGs from organ donors (*5*), we sought to better understand underlying processes and cell types associated with this neuronal loss in DPN hDRGs. To do this, we first subclustered all 8 cell types (**Supplementary Fig. 1A**) to focus on specific subtypes of cells changing from control to diabetes and diabetes to DPN. We conducted Augur cell type prioritization analysis (*7*) identify cell types that become more transcriptionally separable in response to clinical state from control to diabetes and in diabetes to DPN. In the control vs. diabetes condition, we found specific subtypes of Schwann cells, neurons, fibroblasts and adipocytes were prioritized as the most responsive to the clinical diagnosis of diabetes (**Fig. 2A**). Some of these same cell types were again represented in the diabetes to DPN analysis but the single adipocyte cluster now represented the highest prioritization as responsive to the clinical condition followed by a Schwann cell cluster, 2 fibroblast clusters, and 4 subclusters of neurons (**Fig. 2B**). While only 37 genes were differentially expressed in this adipocyte population across clinical conditions (**Supplementary Data 2**), the adipocytes expressed *CIDEA* and *ADIPOQ* suggesting that these cells represent a specific subtype of adipocyte expressing adiponectin and with a metabolic profile influenced by insulin sensitivity (*8*).

**Figure 2.**
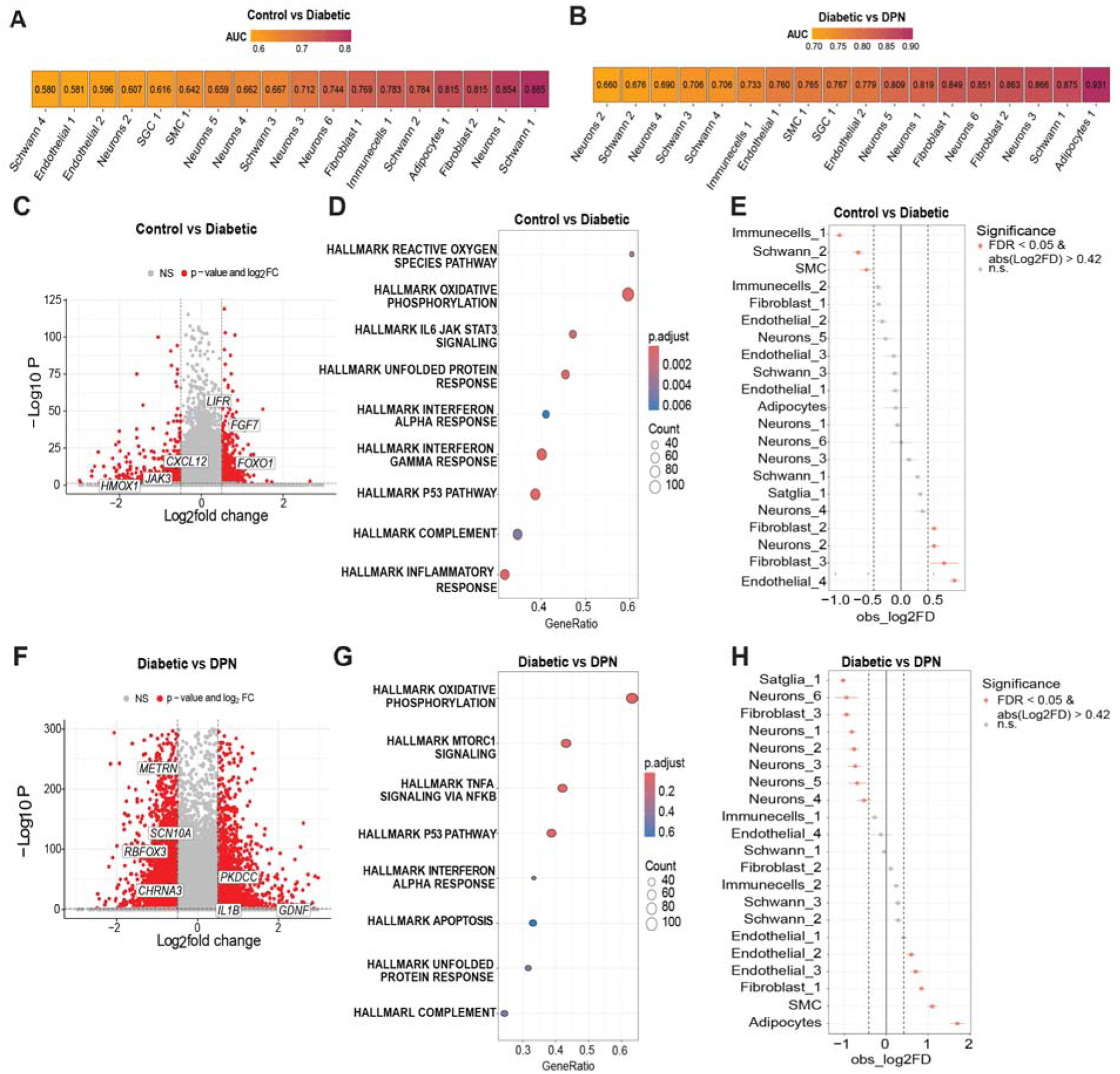
Revealing cell type contributions and transcriptomic alterations in the progression to DPN in the hDRG. (**A–B**) Augur prioritization scores (AUC) for Control vs Diabetic (A) and Diabetic vs DPN (B) comparisons, with cell types ranked by decreasing classification performance; higher AUC values indicate greater transcriptional separability between conditions within that cell type. (**C**) Volcano plot of differentially expressed genes (DEGs) for the Control vs Diabetic comparison across cell types. p-adjusted < 0.05; |log2 fold change| > 0.585. (**D**) Gene set enrichment analysis (GSEA) of DEGs from the Control vs Diabetic comparison, summarizing significantly enriched pathways across cell types (FDR < 0.05); positive normalized enrichment scores (NES) indicate pathways upregulated in Diabetic samples, whereas negative NES indicate pathways downregulated in Diabetic samples. (**E**) Differential subcluster proportions between Control and Diabetic donors, assessed using scProportionTest (Benjamini–Hochberg–adjusted P values; FDR < 0.05; |log2 fold change| > 0.42). (**F**) Volcano plot of DEGs for the Diabetes vs DPN comparison across cell types. p-adjusted < 0.05; |log2 fold change| > 0.585. (**G**) GSEA of DEGs from the Diabetic vs DPN comparison, summarizing significantly enriched pathways across cell types (FDR < 0.05); positive NES indicate pathways upregulated in DPN samples, whereas negative NES indicate pathways downregulated in DPN samples. (**H**) Differential subcluster proportions between Diabetic and DPN donors, assessed using scProportionTest (Benjamini–Hochberg–adjusted P values; FDR < 0.05; |log2 fold change| > 0.585).

Augur analysis provides insight into cell type changes in response to perturbation, in this case clinical diagnosis. To assess differential gene expressions, we pseudo-bulked all cell types by gene and compared control vs. diabetic conditions. In this analysis we found 800 downregulated genes and 688 upregulated genes (**Fig. 2C and Supplementary Data 3**). We then examined the Gene Set Enrichment Analysis (GSEA) Hallmark Genes represented within these differentially expressed genes (*9*). We found enrichment for well-known diabetic neuropathy and neuropathic pain processes like reactive oxygen species and oxidative phosphorylation (*1, 10*), interleukin 6 (IL-6) signaling (*1*), and the unfolded protein response (*11–13*), in addition to other pathways linked to neuropathic pain like the type 1 interferon response (*14, 15*) and complement signaling (*16*). These changes were present even though there was no evidence of neuropathic pain either in family history or medical reports in these organ donors with diabetes suggesting that these changes are not directly related to nociceptive processes but may precede development of neuropathic pain in humans (**Fig. 2D**). We also examined cell type proportions for each subcluster in control vs. diabetes and found decreases in immune cells, a subtype of Schwann cells and smooth muscle cells and increases in 2 fibroblast subtypes, a subtype of endothelial cells and a subtype of neuron in diabetes (**Fig. 2E and Supplementary Fig. 1B**). Comparing diabetic to DPN samples we found 1737 downregulated genes and 1436 upregulated genes in DPN (**Fig. 2F and Supplementary Data 4**). Downregulated genes included *SCN10A*, encoding the voltage-gated sodium channel Nav1.8, which has previously been shown to be downregulated in DRG neurons in preclinical neuropathic pain models (*17*), and the *METRN* gene, encoding meteorin, which plays a key role in pain resolution in preclinical models (*18–22*). Upregulated genes included genes linked to pain signaling, including *PKDCC*, which encodes the vertebrate lonesome kinase (VLK) ectokinase which has recently been identified as key factor linking DRG activity to NMDA receptor clustering at spinal dorsal horn synapses (*23*).

GSEA Hallmark Gene enrichment yielded similar terms with overrepresentation of oxidative phosphorylation, type 1 interferon signaling, unfolded protein response, and p53 pathway signaling, but also revealed altered mechanistic target of rapamycin complex 1 (mTORC1) signaling, tumor necrosis alpha (TNFα) signaling and apoptosis (**Fig. 2G**). Again, many of these pathways have been linked to neuropathic pain in preclinical models, but apoptosis was only represented in the DPN hDRGs. Consistent with the notion that apoptosis might occur in neuronal populations during the transition from diabetes to DPN, 6 board neuronal subclusters were decreased in cell proportion in DPN DRGs while there was an increase in endothelial cells, fibroblasts, smooth muscle cells and adipocytes in DPN DRGs (**Fig. 2H and Supplementary Fig. 1B**).

### Neuronal changes in the progression from diabetes to DPN

To focus on changes in neuronal populations we did label transfer on all of these samples from our recently completed reference atlas for the hDRG (*24*) finding that all 22 neuronal subtypes were represented in our sample (**Fig. 3A and Supplementary Fig. 2**). All neuronal cell types were present in control, diabetic and DPN samples (**Fig. 3B**), however, there were shifts in neuronal cell type proportions seen from one condition to another (**Fig. 3C**). To further understand these population shifts, we did single cell proportion testing on control vs. diabetic and diabetic vs. DPN hDRG samples. In control vs. diabetic, we noted a marked increase in the proportion of ATF3 neurons as well as C-PEP.TAC1/CACNG5 and A-PEP.SCGN/ADRA2C neurons (**Fig. 3D**). Several cell types also showed decreased proportions, and this decrease was consistent in diabetic vs. DPN samples except for A-LTMR.TAC3, C-PEP.TAC1/CACNG5, and C-NP.SST populations which were only found in the diabetic vs. DPN samples (**Fig. 3E**). ATF3, C-NP. MRGPRX1/MRGPRX4, and C-PEP.TAC1/CHRNA3 were increased in the DPN vs. diabetic samples (**Fig. 3E**). Next we examined expression of each of the unique marker genes for neuronal populations identified in the hDRG reference atlas (*24*) and noted a clear increase in expression of genes associated with the ATF3 neuronal cluster (*ATF3, UCN, PENK* and *TRIM54*) from control to diabetes to DPN while many other marker genes showed a trend toward downregulation from control to diabetes to DPN (**Fig. 3F**).

**Figure 3.**
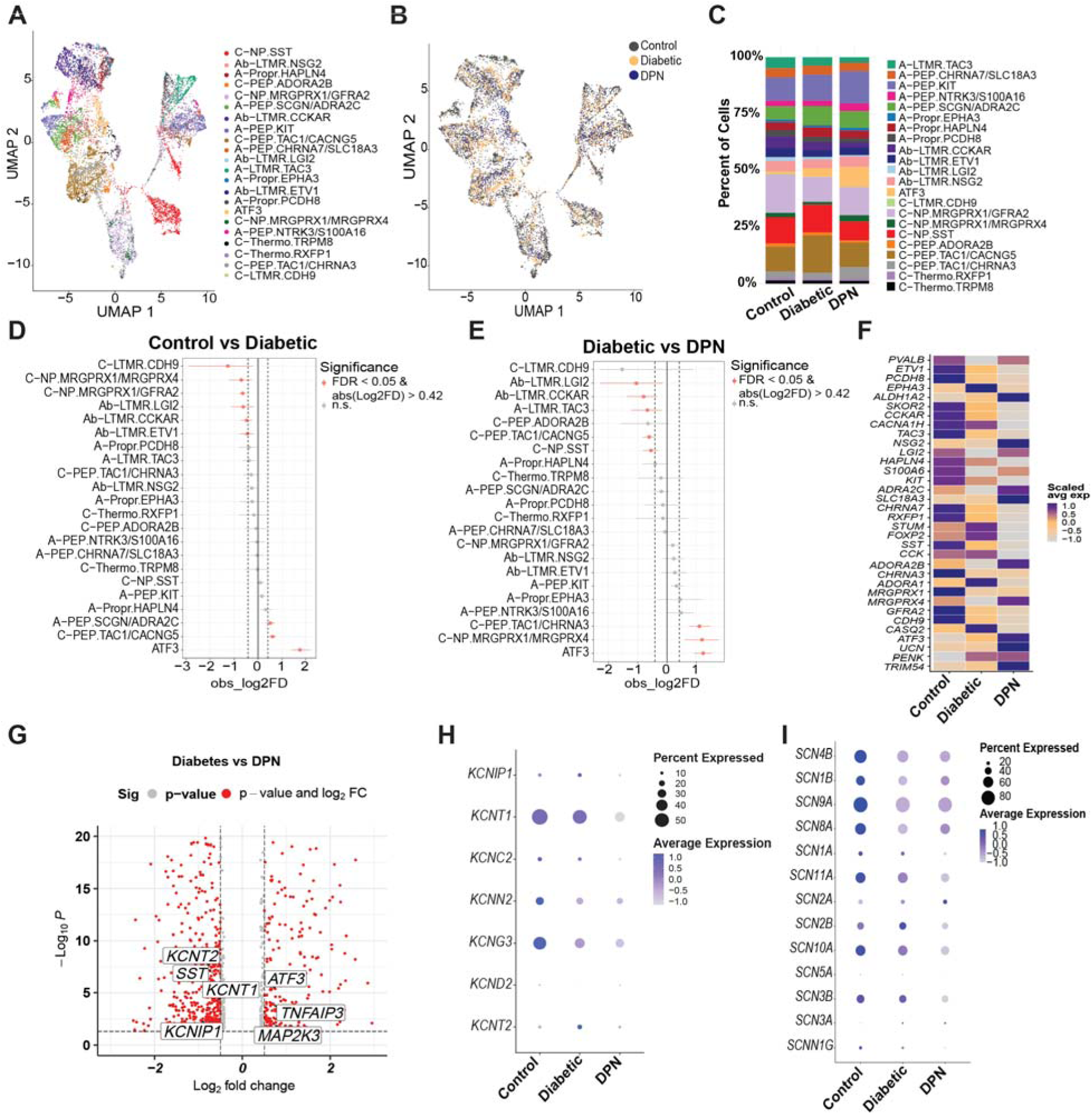
Progressive neurodegeneration of sensory neurons in hDRG accompanies the progression to DPN. (**A**) UMAP projection of single-nucleus RNA-seq (snRNA-seq) neuronal nuclei identifying 22 neuronal subtypes; nuclei are colored by subtype annotation. (**B**) UMAP projections of neuronal nuclei colored by condition (Control, Diabetic, and DPN). (**C**) Relative abundance of neuronal subtypes across conditions based on snRNA-seq. (**D–E**) Differential neuronal subtype proportions between Control vs Diabetic donors (D) and Diabetic vs DPN donors (E), assessed using scProportionTest (Benjamini–Hochberg–adjusted P value (FDR) < 0.05 and |log2 fold change| > 0.585). (**F**) Heatmap of neuronal subtype–specific marker gene expression across conditions; colors indicate scaled average expression within each condition. (**G**) Volcano plot of differentially expressed genes (DEGs) for the Diabetic vs DPN comparison in neurons. p-adjusted < 0.05; |log2 fold change| > 0.585. (**H**) Dot plot of potassium channel DEG expression across conditions; dot size denotes the fraction of nuclei expressing each gene and color denotes scaled, log-normalized average expression. (**I**) Dot plot of voltage-gated sodium channel gene expression across conditions; dot size denotes the fraction of nuclei expressing each gene and color denotes scaled, log-normalized average expression.

Next, we examined differential expressions of genes between the diabetes and DPN samples with all neuronal nuclei pooled for each condition. We found 448 downregulated genes and 298 upregulated genes (**Fig. 3G and Supplemental Data 5**). Downregulated genes included a substantial number of voltage-gated potassium channel genes (**Fig. 3H**), consistent with previous reports in the preclinical neuropathic pain literature (*25*). Moreover, many voltage-gated sodium channels showed a trend toward decreased expression with the progression of diabetes to DPN (**Fig. 3I**), again consistent with preclinical studies of neuropathic pain (*17*).

### Induction of a neuronal, apoptosis-related transcriptional program in diabetes and DPN

Given our observation of a reduced proportion of neuronal populations in DPN compared to diabetes, and increased expression of apoptosis-related genes with DPN, we more thoroughly assessed gene expression in neurons associated with apoptosis. First we examined gene expression for a core set of apoptosis-related genes (*26*) and found that these genes showed a clear trend toward increased expression in hDRG neurons with progression of disease (**Fig. 4A**). This trend was also represented when we examined the number of cells expressing these apoptosis markers across sample types (**Fig. 4B and Supplementary Fig. 3A**). When we queried which neuronal subtypes demonstrated the highest enrichment of apoptosis-related genes, we found clear trends in the ATF3 and several other A-fiber populations (**Fig. 4C**) many of which were also decreased in neuronal proportion in our previous analysis (**Fig. 3E**).

**Figure 4:**
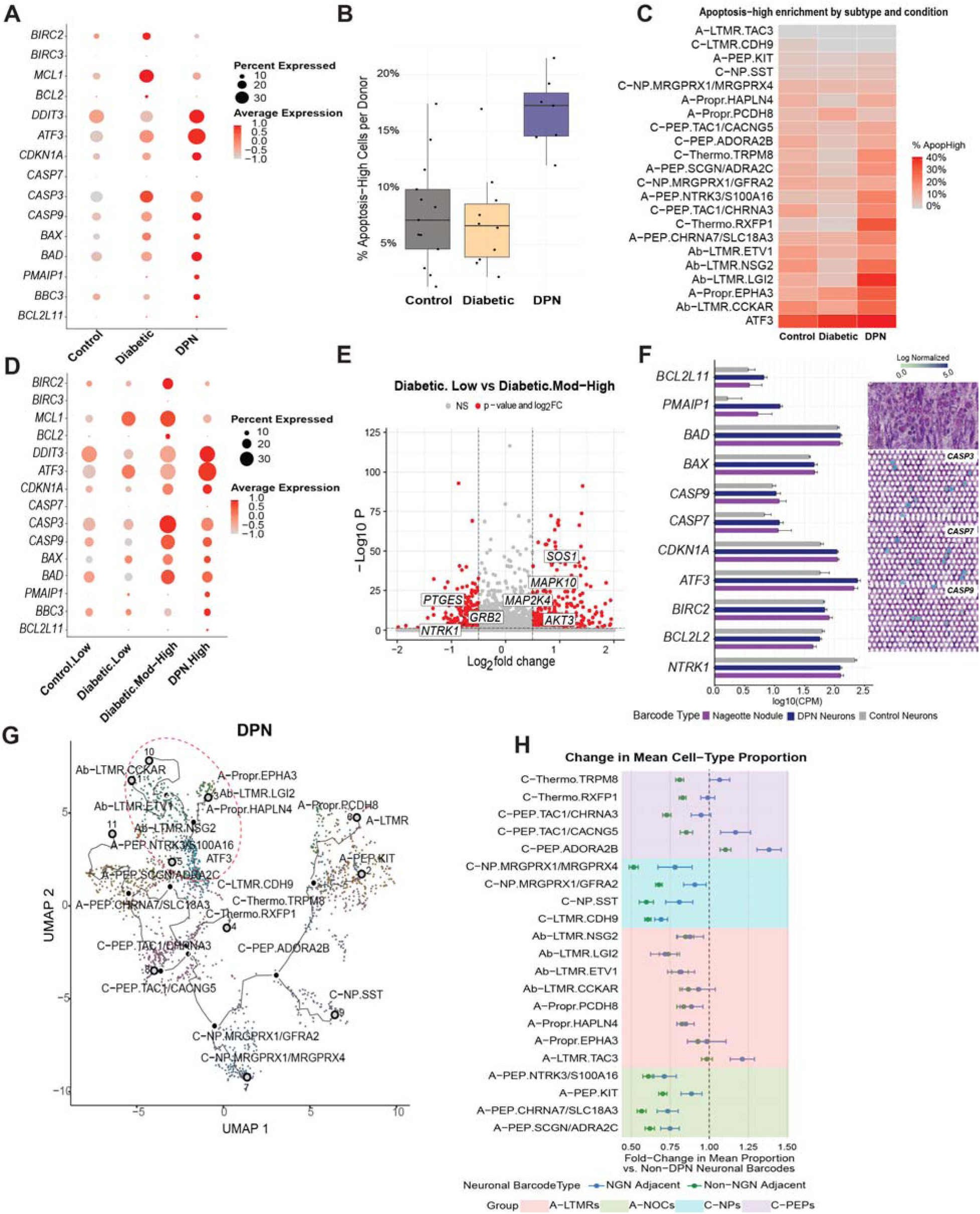
An apoptosis-associated neuronal state increases in hDRG with DPN and is associated with progression of Nageotte nodules. (**A**) Dot plot showing expression of apoptosis- and stress-related genes across conditions (Control, Diabetic, DPN); dot size indicates the percentage of nuclei expressing each gene and color denotes scaled average expression. (**B**) Bar plot showing the percentage of ApopHigh neuronal nuclei per donor across conditions; ApopHigh nuclei are defined as those in the top 10th percentile of the apoptosis score in the dataset. (**C**) Heatmap quantifying ApopHigh enrichment across neuronal subtypes by condition; color indicates the percentage of ApopHigh nuclei within each subtype (Control, Diabetic, DPN). (**D**) Dot plot of apoptosis and stress gene expression stratified by Nageotte score and condition (Control-Low, Diabetic-Low, Diabetic-Mod-High, DPN-High), displayed as percent expressed (dot size) and scaled average expression. (**E**) Volcano plot of differential expression between Diabetic-Low and Diabetic-Mod-High neuronal barcodes, highlighting selected signaling and stress-associated genes. p-adjusted < 0.05; |log2 fold change| > 0.585 (**F**) Left, expression of representative apoptosis/stress genes (counts per million, CPM) across neuronal barcode types (Nageotte nodule–associated, DPN neurons, Control neurons). Right, Visium spatial transcriptomics maps illustrating caspase transcript barcodes (*CASP3/CASP7/CASP9*) localized to neurons. (**G**) Trajectory projections over the neuronal UMAP in DPN; dashed outlines highlight A-fiber subpopulations that project into the ATF3 injury-associated region of the trajectory. (**H**) Deconvolution of Visium barcodes estimating neuronal subtype composition near Nageotte nodules; fold change compares nodule-adjacent versus non-adjacent DPN barcodes (relative to non-DPN barcodes), with shading grouping subtypes into broader classes (A-LTMRs, A-NOCs, C-NPs, C-PEPs).

We then sought to understand whether neuronal apoptosis in the hDRG correlated with the degree to which Nageotte nodules are observed in the samples. To do this, we used our previously developed Nageotte nodule scoring system (*5*) with histological examination of DRG sections from each of the organ donors used in this study. We again examined expression for a core set of apoptosis-related genes and found positive correlation between Nageotte nodule score and gene expression for each gene (**Fig. 4D**). To focus on changes in gene expression that might be associated with a progression towards apoptosis we examined differential gene expression in neuronal nuclei between the low Nageotte nodule samples and the moderate-to-high samples (score 3-4), excluding the high Nageotte nodule (score 4-5) samples that have already undergone severe neurodegeneration. Here we found 153 downregulated genes, including the neuronal survival factor *NTRK1*, which encodes the TrkA receptor for nerve growth factor, and 356 upregulated genes (**Fig. 4E and Supplementary Data 6**). Upregulated genes were enriched for mitogen activated protein kinase (MAPK) genes that are upstream of apoptosis signaling, including the *MAPK10* gene, encoding JNK3 which is a pro-apoptotic signaling factor in ischemic neuronal death (*27*). To provide an independent validation of these findings, we used our previously generated Visium spatial transcriptomic dataset (*5*) to examine expression of a subset of these genes in neuronal barcodes. Here we observed a similar pattern of apoptotic gene expression, markedly increased *ATF3* expression, and decreased expression of the *NTRK1* gene (**Fig. 4F**).

The above analysis raises the question of which neuronal populations turn on apoptotic-related gene expression in diabetes or DPN. To address this open question, we used Monocle3 analysis (*28*) to examine neuronal cell trajectories in each condition. While the control and diabetic trajectory connections were similar, the DPN trajectory analysis demonstrated a shift of the ATF3 neuronal population to a group of A fiber LTMRs suggesting that these neurons become more transcriptomically similar to the ATF3 neuronal population in DPN (**Fig. 4G and Supplementary Fig. 3B**). Importantly, this population of neurons showing similarity of ATF3 neurons was the same group of neuronal populations showing increased apoptosis-related gene expression in **Fig. 4C** (Ab-LTMR.ETV1, Ab-LTMR.NSG2, Ab-LTMR.LGI2, Ab-LTMR.CCKR, A-Propr.EPHA3). This strongly supports the conclusion that Ab-LTMR populations, in particular, are the most susceptible to induction of apoptotic signaling in DPN.

Another unaddressed question raised from our previous work on Nageotte nodules in DPN hDRGs is what neuronal populations are closest to these pathological structures and most likely to sprout into the nodule forming the Nageotte nest structure (*5*). To gain insight into this we did deconvolution analysis of neuronal and Nageotte barcodes in previously published Visium spatial transcriptomics experiments (*5*) and found that two populations of C-fiber neurons were enriched at locations near Nageotte nodules in DPN hDRGs, the C-PEP.TAC1/CACNG5 and C-PEP.ADORA2B populations (**Fig. 4H, Supplemental Fig. 4, and Supplementary Data 7**). While further experiments are needed, these neuronal populations are candidates for neurons that may sprout and form Nageotte nests that we hypothesize could be sites of ectopic electrical activity in DPN hDRGs.

### Computational predictions of ligand-receptor interactions in DPN

To make predictions of potentially important cell-to-neuron interactions that might be responsible for DPN disease progression, and/or neuropathic pain, we constructed a ligand-receptor interactome with ligands from non-neuronal cells to receptors expressed by neurons (*29*). We assembled a list of ligands that were differentially expressed in any non-neuronal cell type between control and diabetes or diabetes and DPN (**Supplemental Data 8**) and a list of receptors that were differentially expressed in any neuronal subtype between control and diabetes or diabetes and DPN (**Supplemental Data 9**). Using a ranking system that considered degree of differential expression for both the ligand and receptor we prioritized the top 50 interactions in the hDRG in DPN (**Fig. 5A**). This group of ligand-receptor interactions included a large number of genes for growth factors like glial-derived neurotrophic factor (GDNF), nerve growth factor (NGF), neurturin, and artemin, synaptic adhesion and axonal growth molecules, and cytokines, consistent with known mechanisms of diabetic neuropathic pain (*1*) and also with our previous finding of dramatic axonal sprouting in DPN hDRGs (*30*).

**Figure 5:**
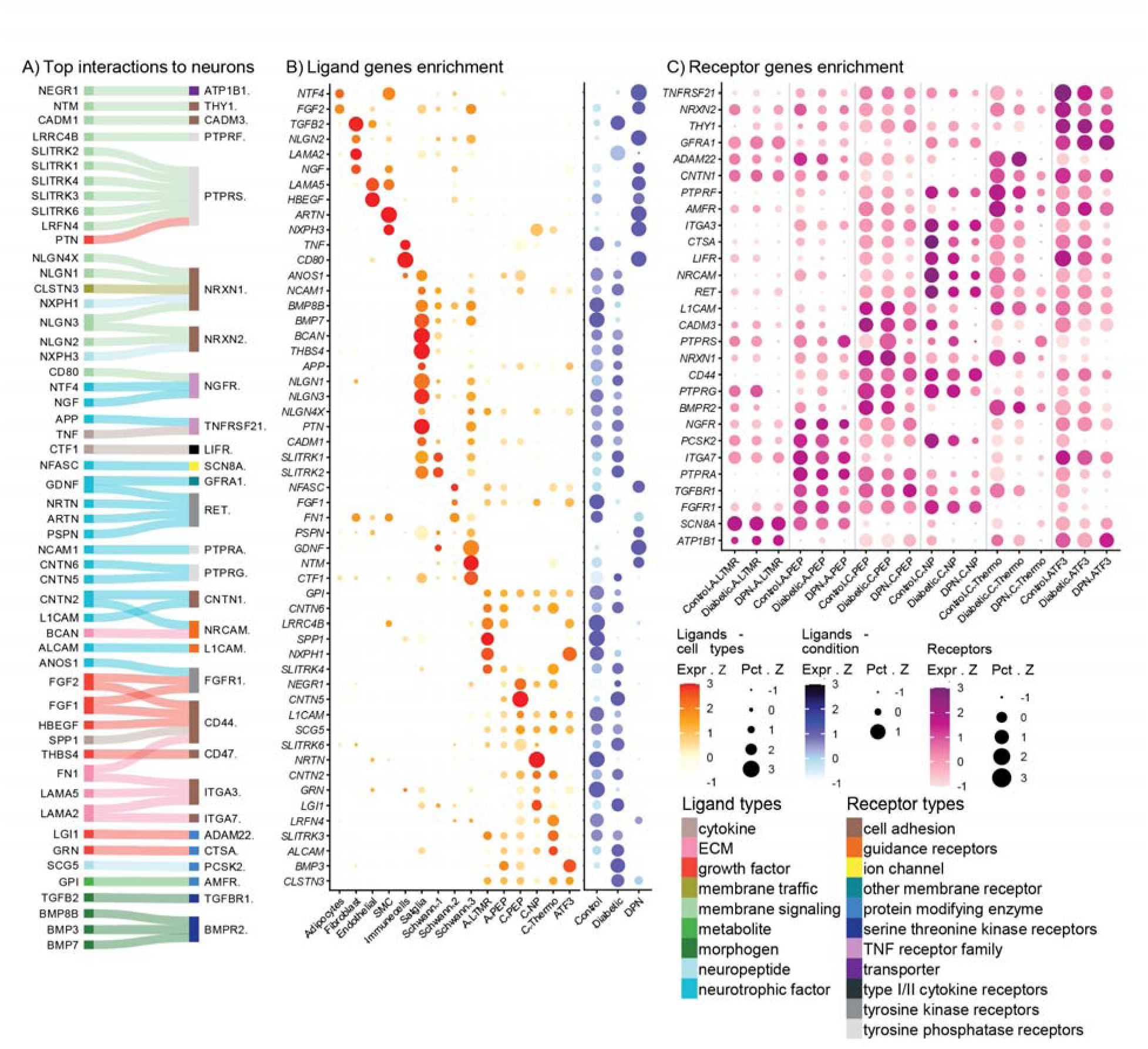
Ligand-receptor interactions in the hDRG potentially driving pathology and pain in the progression to DPN. (**A**) Overview of top differentially expressed ligand–receptor interactions predicted to change with Diabetes and DPN. (**B**) Enrichment of ligand genes across DRG cell types and conditions; dot color represents normalized expression z-scores, and dot size represents the z-score of the percentage of cells expressing each gene. (**C**) Enrichment of receptor genes across neuronal groups by condition; dot color represents normalized expression z-scores, and dot size represents the z-score of the percentage of cells expressing each gene.

To provide insight into the cellular origin of the top 50 ligands shown in **Fig. 5A** we produced dot-plots showing cellular expression, including neurons (while ligand selection was based on differential expression in non-neuronal cells, many of these ligands are also expressed in neurons), for each of these genes (**Fig. 5B**). We also included dot plots showing direction of expression of the gene across cell types from control to diabetes to DPN. The largest number of these ligands originated with either satellite glial cells (SGCs) or with one of 3 subtypes of Schwann cells, although most of the SGC ligands were decreased profoundly in the DPN condition. Interestingly, growth factor genes *NGF, GDNF,* and *ARTN*, were all increased in different cell types with *NGF* in fibroblasts, *GDNF* in Schwann cells, and *ARTN* in endothelial cells, while *NTRN* was decreased and found in endothelial cells and a subtype of neuron (**Fig. 5B**).

We also plotted receptor expression across different subtypes of hDRG neurons grouped by control, diabetic, and DPN conditions. These differentially expressed receptors with ligands in the top 50 differentially expressed list were mostly found in A- or C-fiber nociceptor types and also broadly expressed in ATF3+ neurons (**Fig. 5C**) further supporting the notion that these ligand receptor interactions are important for the neuropathic pain phenotype seen in DPN. This analysis also demonstrates that the biggest changes in expression were seen in ligands from control to diabetes to DPN and not in the receptors expressed by neuronal subtypes.

### Spatial transcriptomics validates differential altered neuronal proportions and gene expression changes in DPN hDRG

We sought to validate findings from single-nuc RNAseq using spatial transcriptomics. To do this, we performed Xenium spatial transcriptomics on 4 control and 4 DPN samples (**Fig. 6A**). We used a Xenium spatial transcriptomics custom gene panel that included cell type markers for neurons and other cell types to enable identification of neuronal cell types from this dataset (**Supplementary Data 10**). Segmentation was performed enabling the identification of 6470 neurons across all samples (**Fig. 6B**) with an average number of transcripts per cell detected of 252 (**Supplementary Fig. 5A**). Using this dataset, we were able to identify all 22 neuronal subtypes described in the reference hDRG atlas (**Fig. 6C and Supplementary Fig. 5B and C**) (*24*). We observed a significant increase in the proportion of the ATF3 neuronal cell type in the DPN group compared to control, validating our single-nuc RNAseq findings (**Fig. 6D and E**). The C-PEP.TAC1/CACNG5 subtype was also increased in DPN hDRGs suggesting that the proximity of Nageotte nodules may shift neurons toward a phenotype consistent with this neuron type (**Fig. 4H**). On the other hand, the C-NP. MRGPRX1/GFRA2 population was significantly decreased in this analysis, which was already decreased from Diabetes to Control in our single-nuc RNAseq analyses (**Supplementary Data 11)**. We also examined differentially expressed genes in our Xenium experiments between control and DPN and found 84 downregulated genes and 38 upregulated genes (**Fig. 6F and Supplementary Data 12**). Many of the genes we identified as downregulated using single-nuc RNAseq were validated in this dataset, including voltage-gated potassium channel genes, *METRN*, and *SCN11A*. Upregulated genes included *OSM, IL1B, CCL3* and *GZMB*, all of which have previously been linked to pain in humans through increased expression in the DRG (*31–33*). In addition, we validated the expression of *PIEZO2, CALCA*, and *P2RX3* using RNAscope and found their expression to be reduced, accompanied by a decrease in the percentage of neurons expressing these genes in samples with DPN (**Supplementary Fig 6**).

**Figure 6:**
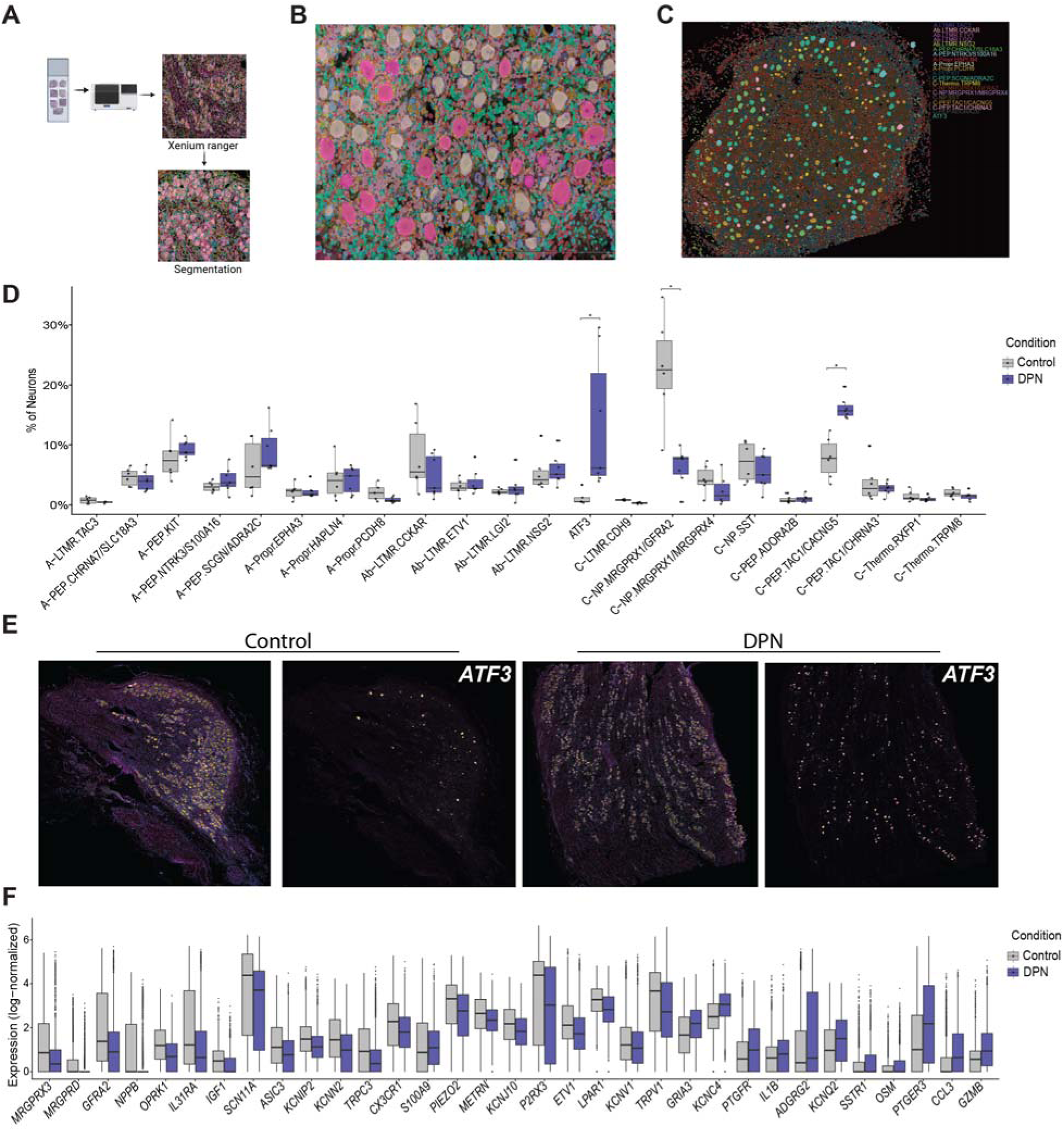
Spatial transcriptomics validates a neuronal neurodegeneration pattern associated with ATF3 expression in the DPN hDRG. (**A**) Overview of the Xenium analysis workflow. (**B**) Representative image showing segmentation of neurons in hDRG. (**C**) Spatial map showing the distribution of annotated neuronal subtypes in hDRG. (**D**) Neuronal subtype composition by condition (Control vs DPN), shown as the percentage of neurons assigned to each subtype per donor (boxplots with individual donors overlaid). Statistical significance was assessed using Kruskal–Wallis tests followed by Dunn’s post hoc test with Benjamini–Hochberg correction for multiple comparisons; p < 0.05 was considered significant. (**G**) Representative Xenium images showing ATF3 transcript signal in Control and DPN sections, highlighting increased expression of the injury-associated marker ATF3 in DPN. (**H**) Boxplots of log-normalized expression for selected differentially expressed genes comparing Control and DPN conditions.

### hDRG bulk proteomics further validates apoptotic signaling in DPN

RNA sequencing provides insight into changes in gene expression but does not always reflect protein expression. To understand changes in the hDRG during the progression of diabetes to DPN, we did data independent-acquisition mass spectrometry (DIA-MS, i.e. quantitative bulk proteomics on 34 control, 9 diabetic, 4 DPN, and 3 non-DPN neuropathic pain organ donors (**Fig. 7A**). In each sample we detected 10,000 and 13,000 proteins and consistently high peptide coverage (**Fig. 7B**). Principle component analysis on all bulk proteomic samples did not reveal any major separation between the groups (**Fig. 7C**) and there were only a small number of differentially expressed proteins (**Fig. 7D and Supplementary Data 13**) likely due to the relatively small sample size and bulk processing of the tissues. Therefore, we sought to examine changes in cellular signaling pathways using GSEA using ranked proteomic changes. Here we found significant overrepresentation of apoptosis, complement and TNFα signaling (**Fig. 7E and Supplementary Data 14**), all consistent with RNAseq findings. Apoptosis was also enriched in DPN samples when accounting for other variables like age, opioid use, and sex of the individual (**Supplementary Fig. 7A**), as well as with CAMERA, a competitive gene set test used as a robust complement to GSEA (**Supplementary Fig. 7B**)To examine whether our proteomic data might represent neuronal loss in the hDRG in DPN we looked at neuronally-enriched proteins and ordered organ donors by disease progression, presence of pain, and Nageotte nodule score.

**Figure 7.**
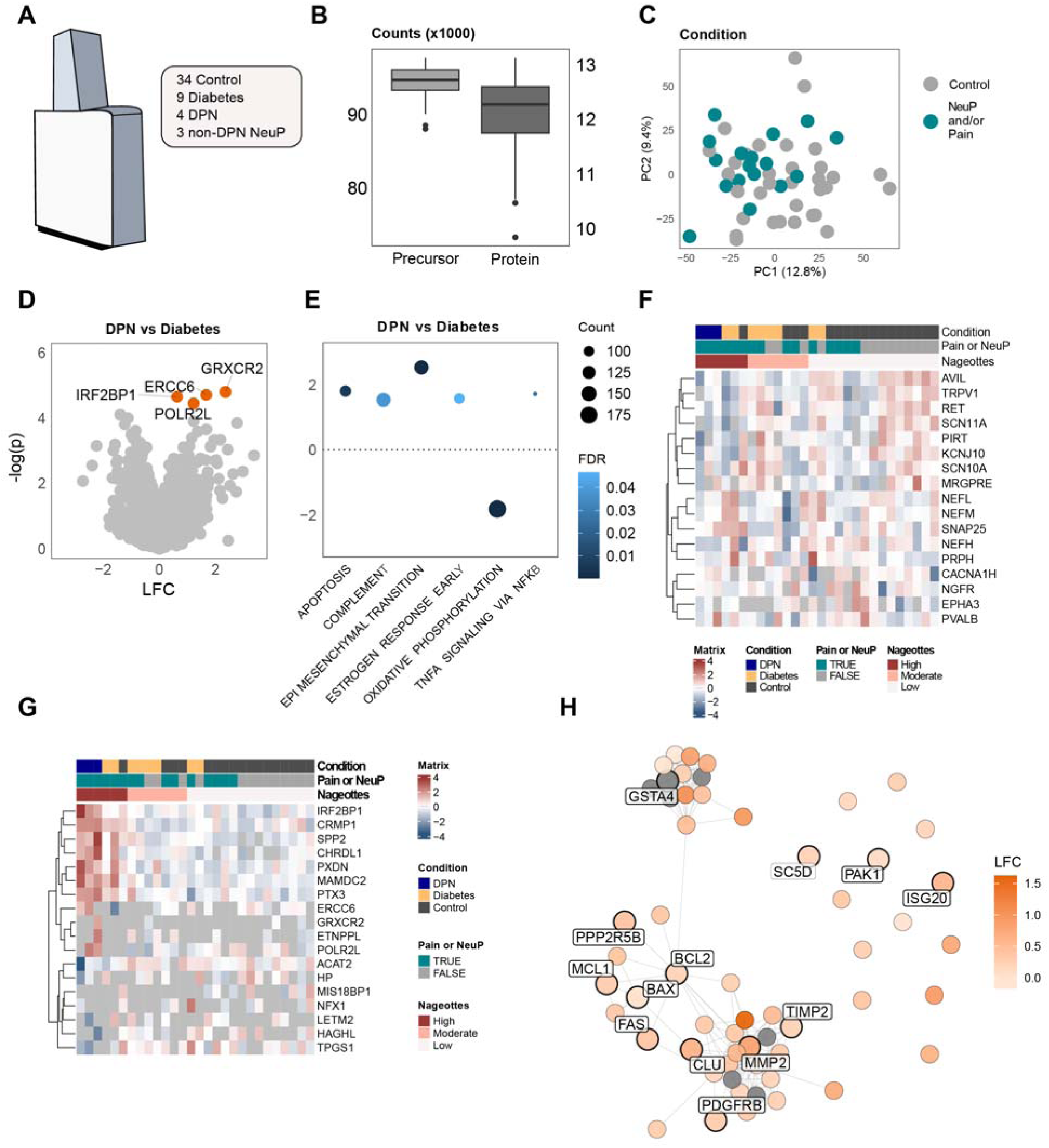
hDRG bulk proteomics validates an apoptosis-associated proteome signature enriched in DPN. (**A**) Schematic overview of the human DRG proteomics cohort, including 34 Control, 9 Diabetic, 4 DPN, and 3 non-DPN NeuP donors. (**B**) Quality control summary showing the number of detected precursors and proteins per sample (counts in thousands). (**C**) Principal component analysis (PCA) of global protein abundance, with samples colored by neuropathy and/or pain status. (**D**) Volcano plot of differential protein abundance for DPN vs Diabetic; proteins meeting FDR < 0.1 are highlighted. (**E**) Gene set enrichment analysis (GSEA) using ranked DPN vs Diabetic proteomic changes; dot color indicates FDR and dot size reflects the number of proteins contributing to each pathway. (**F**) Heatmap of representative neuronal proteins/markers across donors. Values are displayed as row-scaled (z-scored) normalized abundance, with column annotations indicating condition, pain/NeuP status, and Nageotte nodule burden (low/moderate/high). Only donors with associated Nageotte scores are shown.(**G**) Heatmap of representative differentially expressed proteins between Control, Diabetic and DPN donors(Control vs Diabetes, Diabetes vs DPN, Control vs DPN). Values are displayed as row-scaled (z-scored) normalized abundance, with column annotations indicating condition, pain/NeuP status, and Nageotte nodule burden (low/moderate/high). (**G**) STRING protein–protein interaction network of apoptosis- and stress-associated proteins, colored by DPN vs Diabetic log fold change. Nodes denote proteins in the significantly regulated apoptotic pathway, with highlighted nodes also differentially expressed in the snRNA-seq pseudobulk analysis (DESeq2).

Here, only donors with a corresponding Nageotte Scores were plotted. We found a clear underrepresentation of neuronal proteins in the donor samples associated with DPN, neuropathic pain, and high Nageotte nodule score (**Fig. 7F**) and also identified proteins that were significantly regulated across conditions (**Fig. 7G and Supplementary data 15-16**). Apoptosis enrichment was also associated with higher Nageotte nodule scores in the proteomic samples when considering only donors with Diabetes (**Supplementary Fig. 7C**) consistent with transcriptomic findings (**Fig. 4E**). When we examined proteins with GSEA terms associated with apoptosis we found enrichment in the samples with DPN, neuropathic pain, and high Nageotte nodule scores (**Supplementary Fig. 7D and Supplementary data 17-18**). Finally, we conducted STRING protein-protein interaction network analysis examining apoptosis and stress-associated proteins in the hallmark apoptsis pathway uses ranked log-2 fold change (**Fig. 7H**).

Here we found that nodes in the interaction network were consistently composed of proteins also increased as genes in the single-nuc RNAseq of DPN hDRG neurons. This demonstrates that proteomic findings were consistent with RNAseq analysis and highlights a key role for apoptotic signaling in DPN in human sensory neurons.

## Discussion

The main finding emerging from this work is a pattern of neuronal apoptosis signaling-driven neurodegeneration in the hDRG that likely begins early in diabetes becoming severe in DPN. This process is likely irreversible so a key area for future work will be identifying an intervention window for neuroprotection in individuals where diabetes progresses to DPN. We are not aware of previous preclinical studies that find neurodegeneration in the DRG of rodents in any kind of diabetic neuropathy model, but neuronal subtype-biased apoptosis has previously been reported in other nerve injury models (*34*)This may reflect the relatively short duration of diabetes that is generally experimentally induced in these models. Donors obtained in this study frequently had long histories of diabetes and/or DPN prior to death.

We observed signs that A-fiber LTMR and proprioceptive neurons degenerate in individuals with DPN, although neurodegeneration was not exclusive to these kinds of neurons, with a reduced C-NP proportion already in diabetic samples. While an explanation for this selective effect on LTMR and proprioceptive neurons was not tested in our experiments, a recent finding that C-fiber nociceptors are protected from calcium-driven excitotoxicity may yield clues into this effect (*35*). C-fibers express high levels of the non-selective cation channel TRPV1 which allows for large calcium influx when the channel is activated (*36, 37*). TRPV1-expressing neurons display a transcriptomic profile that makes their mitochondria better able to handle calcium overload and protect C-fiber nociceptors from excitotoxicity (*35*). Similar mechanisms may be at play during the progression from diabetes to DPN where two of the best characterized pathologies are increased oxidative stress (*1, 10*), which we also identified in our dataset in this cohort, and mitochondrial dysfunction (*1, 4*), which is usually measured with electron microscopy or mitochondrial signaling assays. A hypothesis emerging from our findings is that neuroprotective strategies should focus on early pathology in A-fiber LTMRs and proprioceptors to prevent degeneration of these neurons, and potentially block further pathological changes like Nageotte nodule formation that occurs after neuronal death (*5*).

Our results encourage a reclassification of DPN as a neurodegenerative disease beyond what is already known about the axonal die back from the epidermis that is often used to diagnose the disease (*3, 4*). Our findings are consistent with the common clinical presentation of DPN where early changes in proprioception and light touch are frequently followed by an insensate and painful neuropathy (*4*). We encourage replication of these findings in larger cohorts development of diagnostic tests that can detect DRG neurodegeneration, and earlier neuroprotective treatment to prevent DPN progression.

## Methods

### Ethical compliance and declaration

All human tissue collection and associated ethical procedures were conducted under Institutional Review Board approval from the University of Texas at Dallas (protocol Legacy-MR-15-237). Dorsal root ganglia (DRGs) were acquired from organ donors through a partnership with the Southwest Transplant Alliance, a Texas-based organ procurement organization. Informed consent for research use was obtained either directly from the donor (first-person authorization) or from a legally authorized representative. Donor screening and consent procedures adhered to United Network for Organ Sharing (UNOS) standards. De-identified clinical information related to each donor was provided by the Southwest Transplant Alliance in accordance with HIPAA regulations to ensure participant privacy.

DRG samples were obtained from 82 human donors (39 female, 43 male) spanning a diverse racial and ethnic background, including 53 White, 13 Hispanic, 15 Black, and 1 donor of unspecified race. Donors were stratified into three clinical groups: (i) healthy organ donors; (ii) donors with diabetes but without a documented diagnosis of clinical neuropathy (some of whom may have been treated with analgesics despite the absence of a clearly recorded neuropathy diagnosis); and (iii) donors with clinically diagnosed diabetic peripheral neuropathy (DPN). For single-nucleus RNA-seq, samples from 14 healthy donors, 10 diabetic donors, and 6 donors with DPN were analyzed; for Visium spatial transcriptomics, DRGs from 3 healthy and 3 DPN donors were included; for Xenium spatial transcriptomics, 4 healthy and 4 DPN samples were profiled; and for proteomics, DRGs from 34 non-diabetic donors, 9 diabetic donors, and 4 donors with DPN were examined.

### Single-nucleus RNA sequencing

Lumbar dorsal root ganglia (DRGs) were collected immediately after recovery and processed without delay to preserve RNA integrity. The dissected DRG tissues were snap-frozen on pulverized dry ice and stored at −80 °C until nuclei extraction. For isolation, frozen tissues were transferred to ice and minced into small fragments (∼1 mm in size) using sterile scalpel blades in ice-cold isolation buffer. The buffer consisted of 0.25 M sucrose, 150 mM KCl, 5 mM MgCl, and 10 mM Tris-HCl (pH 8.0), supplemented with 0.1 mM DTT, a protease inhibitor cocktail, 0.1% Triton X-100, and RNase inhibitor (0.2 U/µL).

The tissue fragments were gently homogenized on ice using a pre-chilled Dounce homogenizer to mechanically dissociate the tissue while preserving nuclear integrity. The homogenate was then filtered through a 100 µm nylon mesh strainer to remove large debris and unbroken tissue. The resulting nuclear suspension was centrifuged at 500 × g for 10 minutes at 4 °C to pellet the nuclei.

The nuclear pellet was gently resuspended in 2 mL of phosphate-buffered saline (PBS) containing 1% bovine serum albumin (BSA) and RNase inhibitor (0.2 U/µL) to stabilize nuclei and protect RNA. A second centrifugation was performed at 500 × g for 5 minutes at 4 °C to further purify the nuclei. The clean nuclear pellet was immediately fixed using the 10x Genomics Chromium Fixed RNA Profiling Kit, according to the manufacturer’s protocol. Fixation was performed at 4 °C for 16 hours to preserve RNA content and nuclear morphology.

Following fixation, nuclei were labeled using the 10x Genomics Fixed RNA Feature Barcode Kit for 16 hours at 4 °C to enable multiplexed assay compatibility. Libraries were constructed from the fixed, barcoded nuclei using the workflow provided by 10x Genomics according to the manufacturer’s instructions(*38*).

Libraries were sequenced in multiple batches using either the Illumina NextSeq 2000 platform at the University of Texas at Dallas Genome Core or the NovaSeq X system at Psomogen.

### Bioinformatic Analysis

Bioinformatic and statistical analyses were conducted in R (v4.5.2). Single-nucleus RNA-seq data were processed and analyzed primarily with Seurat (v5.3.0), together with dplyr (v1.1.4) and the tidyverse (v2.0.0), ggplot2 (v4.0.0 and v4.0.1) for data visualization. ScProportion Test (v0.0.0.9000) was used to identify differences in cell-type proportions between conditions. Gene set annotations were obtained using msigdbr (v25.1.1), and pathway enrichment analyses were performed with clusterProfiler (v4.16.0) and enrichplot (v1.28.4). Pseudobulk differential expression was carried out using DESeq2 (v1.48.0), while single-nucleus–level differential testing used the MAST framework (v1.33.0), as implemented within Seurat. Volcano plots were generated using EnhancedVolcano (v1.26.0), and cell type prioritization was performed with Augur (v1.0.3). Unless otherwise specified, analyses were performed on log-normalized counts from the RNA assay, and multiple hypothesis testing was controlled using the Benjamini–Hochberg procedure to estimate false discovery rate (FDR). For deconvolution analysis python (v3.10.19), scanpy (v1.11.5), numpy(v2.2.6), pandas (v2.3.3), matplotlib(v3.10.7), seaborn(v0.13.2), torch(2.6.0 + CUDA 12.4), and cell2location (v0.1.5) with the parameters N_cells per location = 15, detection alpha= 20 and the q05_cell_abundance_w_sf was selected for cell abundance estimates.

Raw sequencing reads were processed and aligned to the human reference genome (GRCh38) using Cell Ranger v7 (10x Genomics). To eliminate ambient RNA signal SoupX (v1.6.2) was applied with default parameters(*39*).

Integration anchors were performed using canonical correction analysis (CCA) using Seurat v4. Quality control filters were applied to retain 172,740 nuclei with more than 1,000 detected genes and fewer than 10% reads derived from mitochondrial genes. Datawere scaled for downstream analysis. The integrated dataset was visualized and clustered using a shared nearest neighbor (SNN) approach implemented in Seurat’s FindClusters function at a resolution of 0.5, with dimensionality reduction performed via UMAP. Clusters exhibiting enrichment for SNAP25 (log2FC > 0.5, adjusted p < 0.05) were annotated as neurons and canonical markers were used to identify other non-neuronal populations. In addition, label transfer of 22 neuronal subtypes from the reference atlas (*24*) was performed on the neuron object using the label transfer framework implemented in Seurat.

### Cell type prioritization with Augur

To prioritize cell types according to their transcriptional sensitivity to condition, Augur was used as a cell type–prioritization framework (*40*). For each major population, including neuronal subtypes and non-neuronal cells, we first performed sub clustering to define finer cell type subclusters and then applied Augur to determine which subclusters were most responsive to perturbation.

Augur trained a classifier to discriminate between experimental conditions (Control, Diabetic, DPN) based on gene expression profiles and summarized performance using the area under the receiver operating characteristic curve (AUC). Higher AUC values indicate greater transcriptomic separability across conditions and were interpreted as evidence of stronger condition-associated perturbation within a given cell type subcluster. Augur was run on normalized expression matrices using cell type subclusters as the grouping variable and condition as the class label, with default hyperparameters. The resulting AUC scores were compared across cell types, and a heatmap of AUC values was generated using the Heatmap package to visualize the broader landscape of cell type–specific transcriptional responses in Diabetic and DPN samples(*40*).

### Cell type composition and proportional analyses

Changes in cell type composition across conditions were assessed using contingency tables constructed from Seurat metadata. Counts of nuclei per predicted cell type and condition were used to evaluate shifts in cell type abundance. Proportions, including the fraction of each cell type subtype in each condition were visualized as stacked and grouped bar plots. For donor-level proportion comparisons across the three conditions (Control, Diabetic, DPN), Kruskal–Wallis tests were performed, followed by pairwise Wilcoxon rank-sum tests with Benjamini–Hochberg correction for multiple testing where appropriate.

Differences in cell-type proportions between conditions were quantified using the scProportionTest. Analyses were performed on Seurat objects containing sample ID, condition (Control, Diabetic, DPN), and cell-type annotations. For each object, sc_utils was used to build the analysis structure, and permutation test compared proportions between conditions (Control vs Diabetic, Diabetic vs DPN), with cluster identity set to cell type and sample identity to sample ID. An estimated proportion difference, 95% confidence interval, and permutation-based p-value, which were adjusted using the Benjamini–Hochberg procedure. Significant shifts in cell-type proportions were visualized using permutation plot and custom bar plots, emphasizing neuronal subtypes with altered abundance in Diabetic and DPN samples relative to controls(*41*).

### Differential expression and gene set enrichment analysis

Differential expression and gene set enrichment analysis were performed on single-nucleus and pseudobulk data to compare conditions (control to diabetes to DPN). At the single-nucleus level, differential expression (DE) was computed with Seurat’s FindMarkers function using the MAST test on log-normalized expression values, including relevant covariates such as samples(*28*). For pseudobulk analysis, raw counts were aggregated by sample and condition to generate pseudobulk profiles, and DESeq2 (*42*)was applied with a design formula (∼sample +condition) including condition to obtain gene-wise log fold changes and false discovery rate (FDR)–adjusted p-values.

Volcano plots were generated using the EnhancedVolcano package from the differential expression results obtained with Seurat’s FindAllMarkers function. The x-axis represents the Seurat avg_log2FC and the y-axis represents −log10(adjusted p value). Genes with |avg_log2FC|> log (1.33) and FDR < 0.05 were considered significantly differentially expressed and colored red (up- or down-regulated), whereas non-significant genes were colored grey..

For gene set enrichment analysis (GSEA), genes were ranked by log fold change from the DE analyses for each comparison (control to diabetes to DPN), and these ranked gene lists were used as input to the GSEA functions in clusterProfiler, with Hallmark gene sets supplied via msigdbr(*43, 44*). The normalized enrichment scores (NES) and FDR-adjusted p-values were reported and visualized using dotplot and gseaplot2 functions from the enrichplot package.

### Pathway-based apoptosis enrichment score

Apoptotic signaling at single-nucleus resolution was quantified using a composite apoptosis enrichment metric (ApoptosisTriggerScore) computed for each nucleus. Hallmark gene sets were obtained from MSigDB via msigdbr, specifically apoptosis, P53 pathway, DNA repair, and unfolded protein response (UPR), and per-nucleus module scores for each gene set were calculated on log-normalized RNA expression using Seurat’s AddModuleScore function.

To account for opposing pro-survival signaling, an anti-apoptotic “survival” module was constructed from canonical anti-apoptotic and inhibitor-of-apoptosis genes, including BCL2 family members and IAP-related factors (e.g. *BCL2, BCL2L1, BCL2L2, BCL2A1, MCL1, XIAP, BIRC2, BIRC3, BIRC5, BIRC6, CFLAR*) together with NF-κB–linked regulators. The ApoptosisTriggerScore was then defined as ApoptosisTriggerScore = (apoptosis module) + (P53 module) + (DNA repair module) + (UPR module) − (survival module), providing a single continuous measure that integrates apoptotic signaling, DNA damage response, ER stress, and pro-survival tone. Distributions of the ApoptosisTriggerScore across conditions (Control, Diabetic, DPN) were visualized using heatmap(*45, 46*).

To define apoptosis-primed cells, an apoptosis-high (ApopHigh) population was identified based on the distribution of the ApoptosisTriggerScore across all neurons. The 90th percentile of the ApoptosisTriggerScore was computed, and nuclei with scores at or above this threshold were labeled ApopHigh, with the remaining nuclei classified as ApopLow (*47*). For each condition and, where indicated, for each neuronal subtype, the proportion of ApopHigh nuclei among all neurons was calculated and used to quantify enrichment of apoptosis-primed cells in Diabetic and DPN samples relative to Control.

Cell type labels were taken from the neuronal sub populations annotationed in the Seurat object. For each combination of neuronal subtype and condition (the total number of nuclei, the number of ApopHigh nuclei, and the fraction of ApopHigh nuclei (PropHigh = ApopHigh / total) were computed. These summaries were used to identify neuronal subtypes showing disproportionate enrichment of apoptosis-high nuclei across conditions, and results were visualized using barplots with condition on the x-axis and the percentage of ApopHigh nuclei on the y-axis.

To account for donor-level variability, donor-resolved analyses were also performed. For each donor and condition, the proportion of ApopHigh nuclei was calculated either across all neurons or within specific neuronal subtypes, and these donor-level proportions were compared across the three conditions using Kruskal–Wallis tests, followed, where indicated, by pairwise Wilcoxon rank-sum tests with Benjamini–Hochberg correction for multiple comparisons.

### Trajectory analysis

Trajectory analysis was performed using monocle3 (v1.4.26) on neurons.. The neuronal object was converted to a cell_data_set using SeuratWrappers, and existing UMAP embeddings were used as the low-dimensional space for trajectory inference. A principal graph was then learned across all neurons with a single partition to capture continuous transcriptional transitions between neuronal states. For visualization, cells were projected onto the learned trajectory and colored by neuronal subtype, with branch points and terminal states labeled. Condition-specific trajectories were examined by sub setting based on condition (Control, Diabetic, and DPN) and replotting the shared trajectory graph for each subset to assess alterations in lineage structure across disease states.

### Nageotte scoring scheme

Nageotte scores for human DRGs were assigned using a previously described qualitative 5-point scale (*5*), in which each ganglion is rated from low to high prevalence of Nageotte nodules based on hematoxylin and eosin histopathology. Scores of 1–2 were classified as Low, scores of 3–4 as Moderate–High, and scores of 4–5 as High Nageotte burden. Nageotte scores were available for 47 donors in this study.

### Statistical analysis

Continuous metrics such as the ApoptosisTriggerScore and individual Hallmark module scores typically showed non-Gaussian, skewed distributions, so non-parametric tests were used. For two-group comparisons (e.g. Diabetic vs Control, DPN vs Diabetic), the Wilcoxon rank-sum test was applied; for comparisons across three conditions (Control, Diabetic, DPN), Kruskal–Wallis tests were used. These tests were performed both at the per-cell level (to assess distributional shifts) and at the per-donor level (using donor means or donor-level proportions) to respect donor-level replication. Unless otherwise stated, p-values were adjusted for multiple comparisons using the Benjamini–Hochberg method.

### Spatial deconvolution and analysis

The 10x Visium V1 barcoded spatial transcriptomics assay was used to analyze 16 samples from 6 DRGs (**Table. 1**). The 10x VISIUM protocol was followed exactly as described by the manufacturer (10X Genomics, (*48*)) with a permeabilization time of 12 minutes. The Genome Center in the University of Texas at Dallas Research Core Facilities performed library preparation and sequencing with Illumina Nextseq 2000. Visium frames were imaged with an Olympus vs120 slide scanner. The 10x Space Ranger pipeline (v2.0.0 with 10x reference genome release GRCh38 2020-A/Ensembl98) was run with default parameters to process raw sequencing files and quantify gene counts per barcode. This assay provides near-single neuron resolution, allowing identification and analysis of neuronal subtypes and surrounding cells(*49*).

To prepare a reference for deconvolution, the single-nuclei data was down-sampled to include 50 randomly selected cells per neuronal and non-neuronal cell type per sample, yielding a reference dataset of 19,646 cells. This reference was used to generate a signature matrix with the spatial deconvolution tool cell2location(*50, 51*), which has been shown to effectively differentiate over 50 cell types, including subpopulations with subtle differences, in nervous system tissue(*52*).

Subsequently, cell2location was used to deconvolute the Visium samples and estimate the number of each neuronal and non-neuronal cell type per barcode. The plotting functions described in the cell2location tutorial (cell2location.readthedocs.io/) were used to visualize localization and abundance of cell types in a representative DPN and control sample.

Barcodes containing neurons and Nageotte nodules were also manually annotated using 10x LoupeBrowser for each Visium sample. Barcodes with single neurons were selected for comparison of the proportion of each neuronal population in DPN and non-diabetic Visium samples. The mean proportion in neurons of non-diabetic samples, in neurons of DPN samples adjacent to Nageotte nodules, and in neurons of DPN samples not adjacent to nodules were calculated for each neuron type. The fold-change in proportion with DPN (among nodule-adjacent neurons and non-adjacent neurons) for each neuron type was visualized with ggplot2..

### Interactome analysis

We built an interactome of predicted changes in intra-DRG signaling to neurons with diabetes and DPN using a curated database of ligand-receptor pairs(*54*) and the differential expression results from the single-nuclei data. Ligand genes expressed in the DRG and corresponding receptor genes expressed in neurons were identified by intersecting gene expression data with the ligan-receptor database in the Sensoryomics Interactome web application (sensoryomics.shinyapps.io/Interactome). Ligand-receptor interactions were filtered to include only those that met the following criteria:

Either the ligand gene must be differentially expressed among DRG cells with p-adj. <0.05 (with diabetes compared to no diabetes, with DPN compared to non-neuropathic diabetes, or with DPN compared to no diabetes), or the receptor gene must be differentially expressed among DRG neurons with p-adj. <0.05 in any of the three comparisons. The ligand and receptor gene cannot be differentially expressed in opposite directions for a given comparison.

The receptor gene must be expressed in at least one of the 22 neuronal populations-with expression threshold defined as a log-normalized expression value of at least 0.01 (following application of Seurat’s NormalizeData() on the pseudobulk of each population with default parameters). The receptor must also be expressed in at least 5% of all neurons in one of the conditions (control, diabetes, or DPN). The ligand gene must be expressed in at least 0.1% of all DRG cells in one of the conditions (control, diabetes, or DPN).

A table was compiled of all interactions predicted to change by comparison (control vs diabetes, diabetes vs DPN). There were a total of 1179 interactions across the 2 comparisons with 788 unique interactions. All interactions were ranked by generating a score derived from the log2 fold-changes, adj. p-values, and percent expression values of the ligand and receptor genes. The top 50 unique interactions were visualized with a sankey plot using SankeyMATIC (sankeymatic.com).

For each ligand gene, enrichment among DRG cell types was visualized in a dot plot with color representing the z-scores of normalized expression values and dot size representing z-scores of the percentage of cells expressing the gene. Similarly, a dot plot was generated to show enrichment of ligand genes across all cells in the three conditions, and enrichment of receptor genes in neuronal groups per condition. The scripts used for interactome analysis and figure generation are publicly available on the SPARC portal(*53*).

### Xenium spatial transcriptomics

Xenium in situ gene expression profiling was carried out on fresh-frozen lumbar DRG tissue obtained from 8 samples (4 control and 4 DPN). Tissue was gradually embedded in OCT on dry ice to prevent thawing, cryosectioned at 10 µm directly onto Xenium slides, and stored at −80 °C until processing. Before running the Xenium assay, sections were fixed for 30 min at room temperature, rinsed in PBS, permeabilized in 1% SDS for 2 min, washed in PBS, incubated in 70% methanol on ice for 60 min, and then washed again in PBS.

Following fixation and permeabilization, slides were mounted into Xenium cassettes and processed using the Xenium in Situ Gene Expression workflow (10x Genomics, User Guide CG000582) with a custom 479-gene panel optimized for DRG. The protocol included overnight probe hybridization, ligation of circularizable probes, and enzymatic amplification of gene-specific barcodes, followed by autofluorescence quenching and DAPI staining. Imaging was performed on the Xenium Analyzer and transcripts were subsequently decoded and explored with Xenium Explorer.

Cell segmentation of lumbar DRG Xenium datasets was performed using Xenium Ranger (v3.1.1) with segmenting out large cells for downstream analysis. The segmented cells were imported into R and Seurat pipelines and were performed for integration and further analyses. Neuronal subtype annotations were assigned by transferring cell type labels from a reference single-cell RNA-seq atlas using Seurat’s label transfer workflow (*24*). Xenium data were visualized and exported via Xenium Explorer (v4.0.0).

### RNAscope in situ hybridization

RNAscope in situ hybridization was performed using the RNAscope Multiplex Fluorescent v1 assay according to the manufacturer’s protocol (Advanced Cell Diagnostics). Assay conditions and tissue handling followed previously described procedures for DRG tissue (*55*).

The following probes were used: *PIEZO2* (449951), *P2RX3* (406301), *CALCA* (605551). Manufacturer-supplied positive control probes were included on each run to verify RNA integrity, and a negative control probe was used to assess background signal. Slides were imaged on an Olympus FV4000 confocal microscope using a 20× objective and were processed using cellSens software (v1.18).

### Quantitative Mass Spectrometry

Data-independent acquisition mass spectrometry (DIA-MS) was performed on human DRG from 50 donors at the Bruker Center of Excellence for Metaproteomics (University of Vienna) as previously described (*56*) with updated data analysis. Samples were processed in 2 batches, with 8 donors in batch 1 (previously published) (*57*) and an additional 42 donors here in batch 2 representing a diverse group of individuals with Diabetes and DPN, as well as other mixed pain conditions, opioid use disorder, and pain-free controls across adulthood.

Spectral deconvolution was performed across all samples using DIA-NN (1.9.1, MBR enabled)(*58*). MaxLFQ was calculated using the diann package (R, v1.0.1) and the technical replicates for batch 1 were averaged by donor to generate a protein expression matrix across donors. PCA was performed on proteins expressed in all samples.

Proteins were filtered for those expressed in > 30% of samples, gene name duplicates removed, and expression modelled through limma (v3.64.3) as ‘∼0 + batch + Diabetic.Type + Opioid.Use + Sex’ unless otherwise mentioned (*59*). Where Nageotte burden was considered, data were first subset for donors with available scores before modelling batch and Opioid.Use with a combined factor of Diabetic.Type and Nageotte Score.

Gene set enrichment analyses were performed using the ClusterProfiler (v4.16.0) package against hallmark gene sets (*60*). Sample based permutations were performed with ‘camerà (limma), and a sensitivity analysis was performed across cohort factors by modelling additional terms (eg. Age, Pain). An FDR was set at 0.05 for pathway enrichment and 0.1 for differentially expressed proteins.High confidence (0.7) protein-protein interaction networks were generated in Cytoscape StringApp (v3.10.3) and plotted in R with Rcy3 (v2.28.1) (*61, 62*).

## Data Availability

Processed sequencing datasets will be deposited in public repositories (dbGaP PRIDE andSPARC: (10.26275/vz0c-uet7 and 10.26275/0jnf-7wq9) with proteomic data available on PRIDE, PXD071905), and processed summary data will be accessible through sensoryomics.com. Analysis code will be released via a public GitHub repository.

## Acknowledgments

The authors thank members of the Price laboratory for human DRG recoveries and the genome center at University of Texas at Dallas . The authors thank members of the Southwest Transplant Alliance for coordination of organ donor tissue recoveries. The authors are grateful to the organ donors and their families for their gift. Proteome data acquisition has been achieved at the Bruker Center of Excellence for Metaproteomics (University of Vienna) and data analysis has been achieved using the Austrian Scientific Computing (ASC) infrastructure.

## Funding

National Institutes of Health grant U19NS130608-01 (TJP, MC, and PMD); Austrian Science Fund 10.55776/PAT2329024 and 10.55776/P36554/ (MS)

## Author contributions

Conceptualization: IS, TJP

Methodology: IS, KM, AMB, SS, AW, MAW, EV, PH, AC, TK, GF

Investigation: IS, KM, AMB, SS, AW, NI, MAW, ECM, DT-F, MSY, JRS, FX, MS

Visualization: IS, KM, AMB, ECM, DT-F

Funding acquisition: SS, MSY, DT-F, GD, PMD, MC, MS, TJP

Project administration: PMD, MC, TJP

Supervision: GD, MS, TJP

Writing – original draft: IS, TJP

Writing – review & editing: all authors

## Competing interests

TJP is co-founder of 4E Therapeutics, PARMedics, NuvoNuro, Nerveli, and Ted and Greg’s. MC is chief medical officer of 4E Therapeutics. MS have an ongoing scientific collaboration with Bruker (Bruker Center of Excellence for Metaproteomics, University of Vienna), however, this collaboration did not influence the content of the manuscript. The other authors declare they have no competing interests.

## Notes

### Competing Interest Statement

TJP is co-founder of 4E Therapeutics, PARMedics, NuvoNuro, Nerveli, and Ted and Gregs. MC is chief medical officer of 4E Therapeutics. MS have an ongoing scientific collaboration with Bruker (Bruker Center of Excellence for Metaproteomics, University of Vienna), however, this collaboration did not influence the content of the manuscript. The other authors declare they have no competing interests.

