## Supplemental Figures and Material for "Progressive neurodegeneration in human dorsal root ganglion from diabetes to painful neuropathy"

**Supplementary Material for:**
**Title: Progressive neurodegeneration in human dorsal root ganglion from diabetes to painful neuropathy**

**Authors:** Ishwarya Sankaranarayanan[1,*], Khadijah Mazhar[1], Allison M. Barry[1], Stephanie Shiers[1], Andi Wangzhou[1], Julia R. Sondermann[5], Feng Xian[5], Nikhil Inturi[1], Michael A. Wilde[1], Eric C. Meyers[6], Joseph .B. Lesnak[1], Jayden A. O'Brien[1], Muhammad Saad Yousuf[1], Manuela Schmidt[5], Gregory Dussor[1], Erin Vines[2], Peter Horton, Anna Cervantes, Tariq Khan, Geoffrey Funk, Diana Tavares-Ferreira[1,] PRECISION Human Pain Network, Patrick M. Dougherty[3], Michele Curatolo[4], Theodore J. Price[1,*]

**Supplemental Figures:**

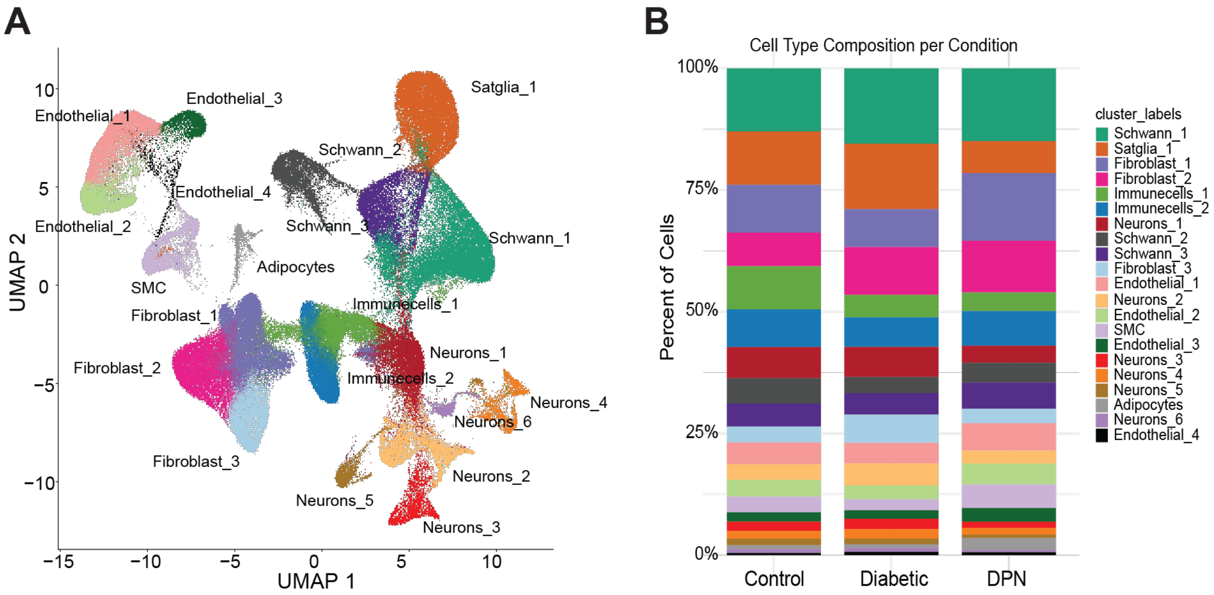


Figure S1

**Supplementary Fig. 1: Single-nucleus RNA-seq overview of hDRG across Conditions.**

(**A**) UMAP projection of hDRG single-nucleus RNA-seq (snRNA-seq) data showing major cell types and associated subclusters.

(**B**) Relative abundance of cell type subclusters across conditions based on snRNA-seq.

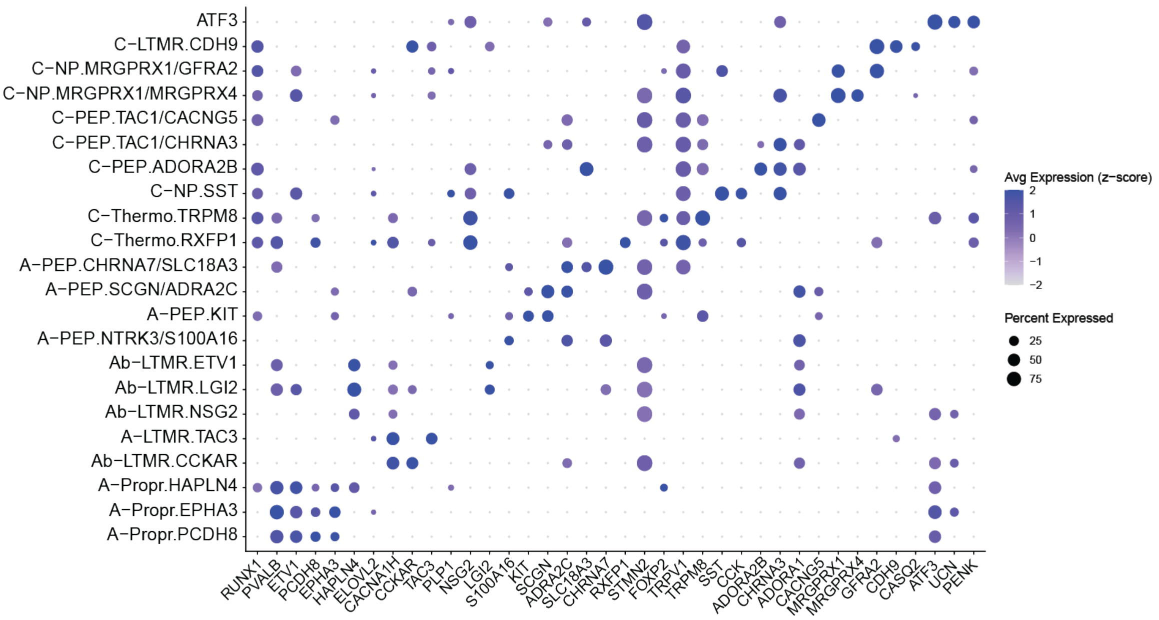


Figure S2

**Supplementary Fig. 2: Cell type specific gene expression of 22 subtypes of Neurons.**

Dot plots of representative marker genes used to annotate major subtypes of neurons. dot color represents normalized expression z-scores, and dot size represents the z-score of the percentage of cells expressing each gene.

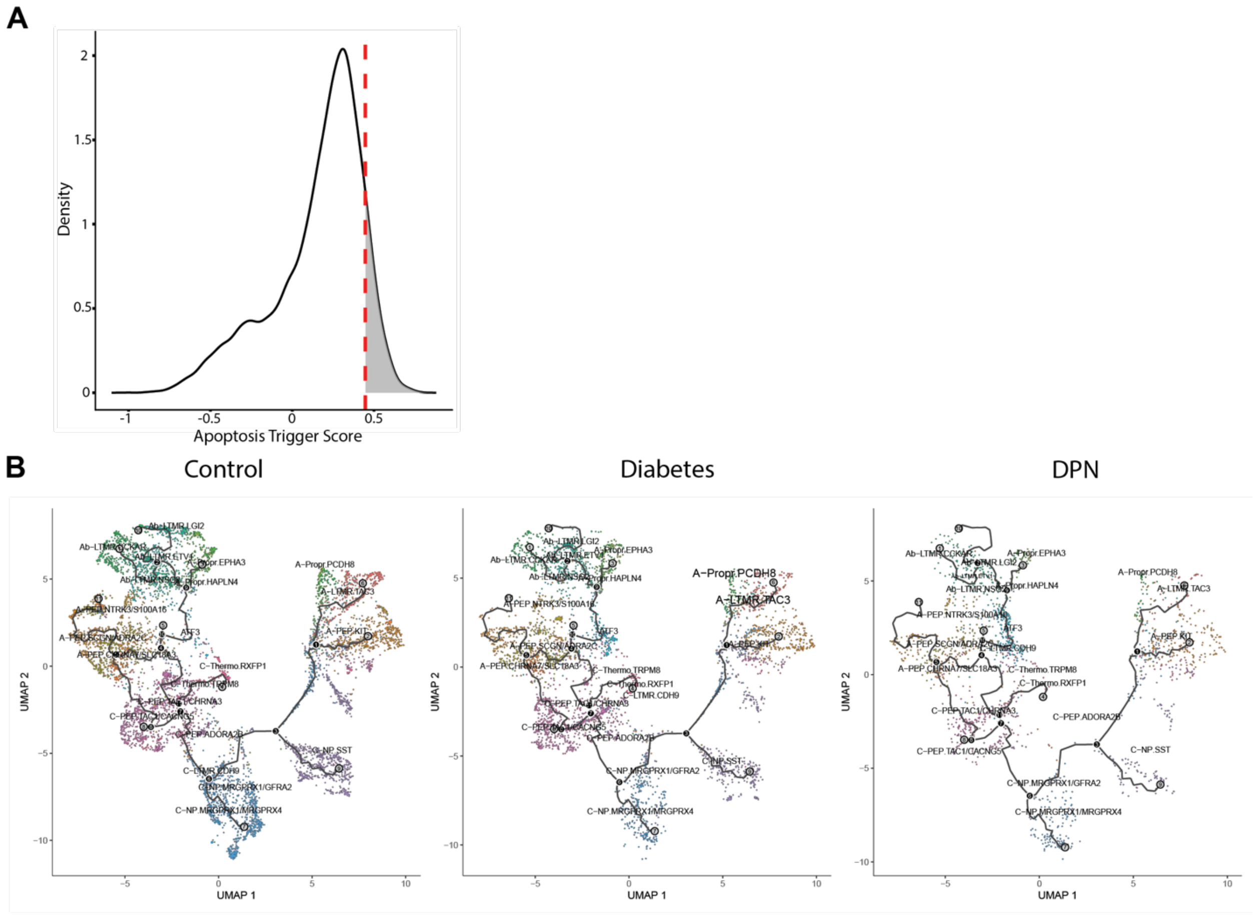


Figure S3

**Supplementary Fig. 3: Trajectory projections of neuronal transcriptional states across Control, Diabetic, and DPN hDRG.**

Trajectory projections over the neuronal UMAP across conditions (Control, Diabetes, DPN)

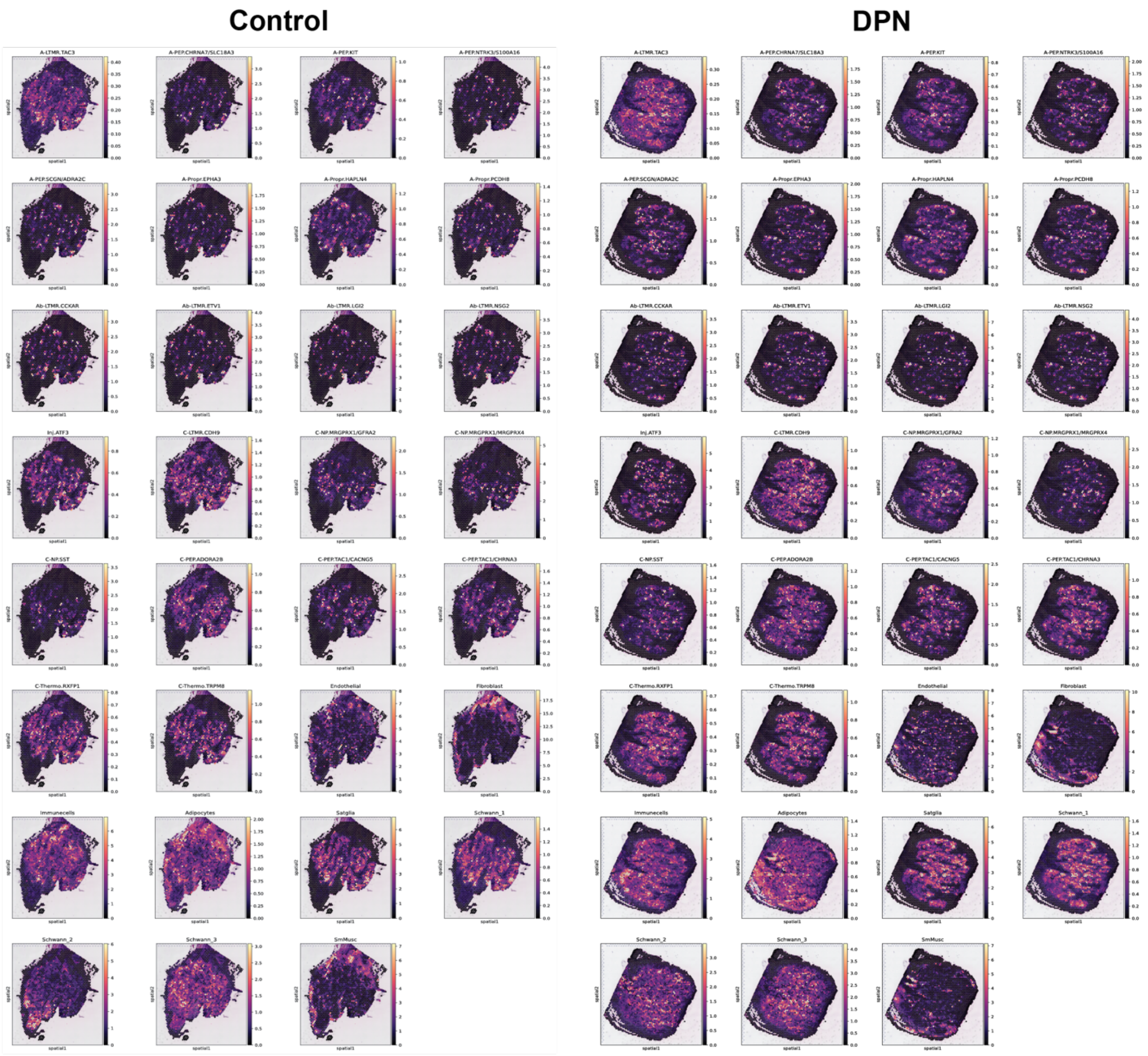


Figure S4

**Supplementary Fig. 4: Deconvolution of Visium barcodes reveals neuronal subtype and non-neuronal cell composition in Control and DPN hDRG.**

**(A)** Density plot of apoptosis trigger scores, with the shaded region indicating cells in the top 10th percentile (apoptosis-high cells).(B) Representative images of deconvolution of Visium barcodes estimating neuronal subtype composition and non-neuronal cell types across Control and DPN conditions.

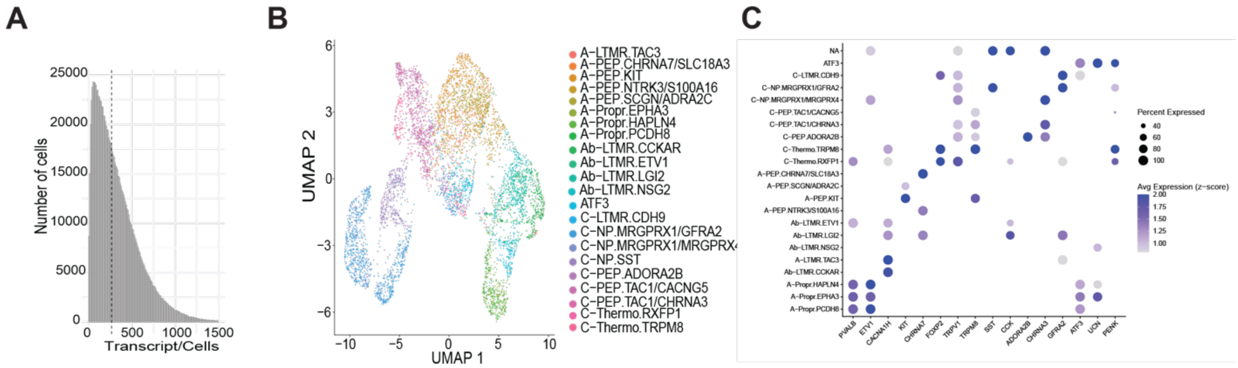


Figure S5

**Supplementary Fig. 5: Xenium in situ profiling and annotation of hDRG neuronal subtypes**

(**A**) Quality-control distribution of total transcripts detected per segmented cell.

(**B**) UMAP projection of neurons using Xenium in situ, colored by neuronal subtype annotation.

(**C**) Dot plots of representative marker genes used to annotate major subtypes of neurons in Xenium. dot color represents normalized expression z-scores, and dot size represents the z-score of the percentage of cells expressing each gene.

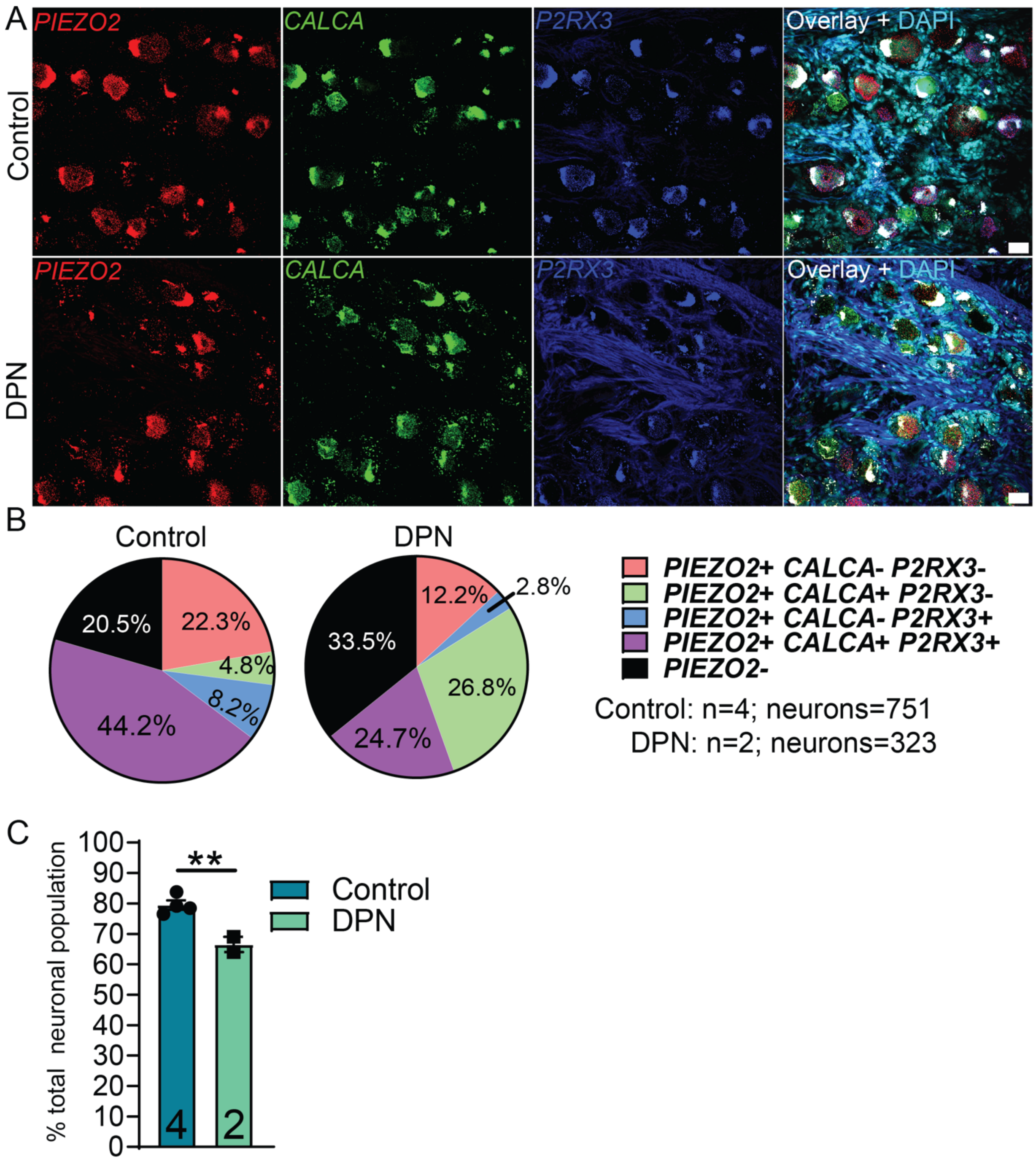
A
PIEZO2
CALCA
P2RX3
Overlay + DAPI
Control
DPN
B
Control
DPN
22.3%
4.8%
8.2%
44.2%
20.5%
12.2%
2.8%
26.8%
24.7%
33.5%
PIEZO2+ CALCA- P2RX3-
PIEZO2+ CALCA+ P2RX3-
PIEZO2+ CALCA- P2RX3+
PIEZO2+ CALCA+ P2RX3+
PIEZO2-
Control: n=4; neurons=751
DPN: n=2; neurons=323
C
% total neuronal population
0
10
20
30
40
50
60
70
80
90
100
**
Control
DPN
4
2

Figure S6

**Supplementary Fig. 6: Validation of PIEZO2 loss in DPN hDRG using RNAscope**

(**A**) Representative RNAscope images showing PIEZO2 (red), CALCA (green), and P2RX3 (blue) in hDRG from Control and DPN donors.

(**B**) Quantification of PIEZO2-, CALCA-, and P2RX3-positive neuronal populations in hDRG from Control and DPN donors.

(**C**) Percentage of neuronal population in hDRG comparing Control and DPN donors.

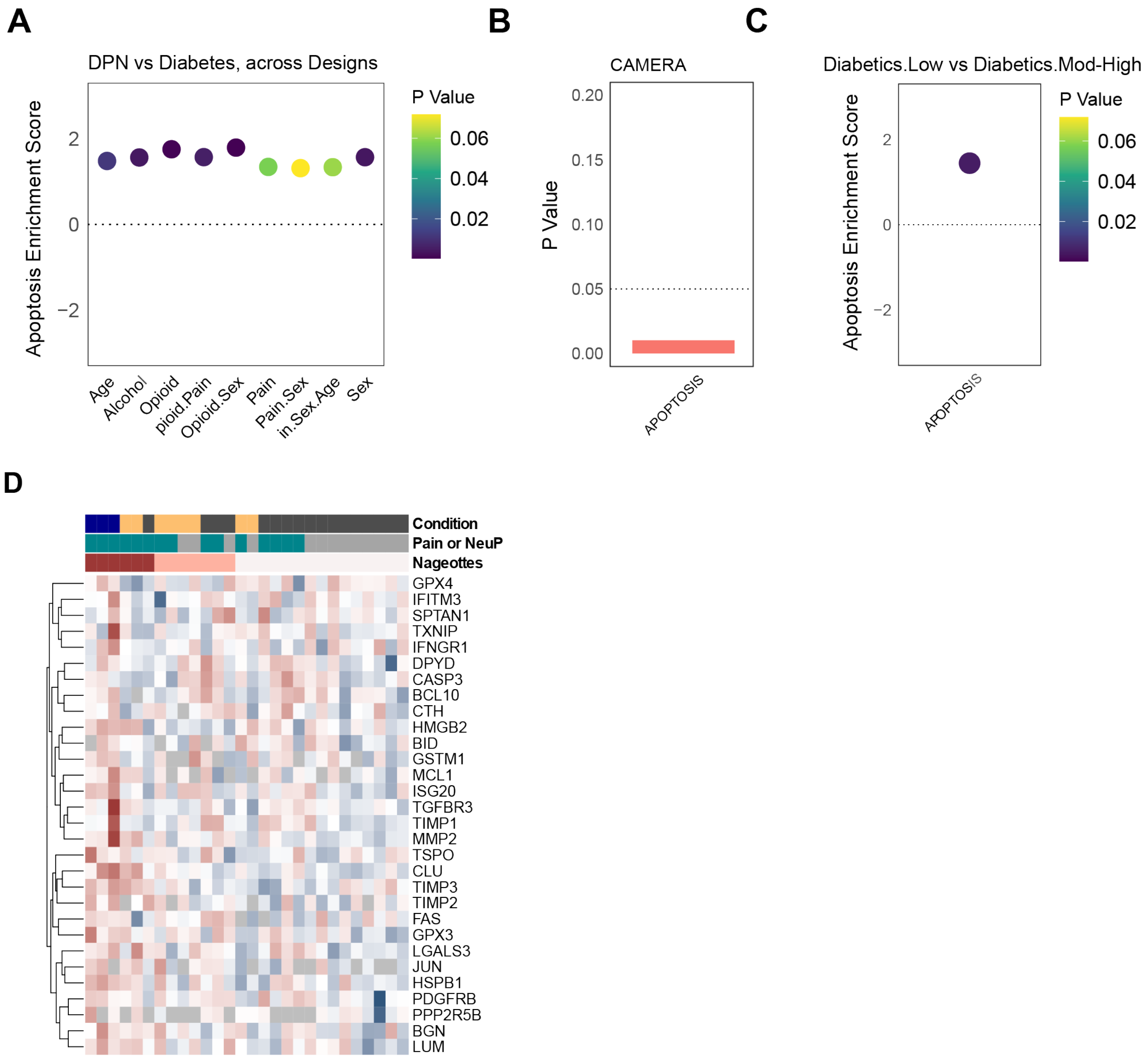


Figure S7

**Supplementary Fig. 7: Apoptosis-associated transcriptomic and proteomic signatures in DPN hDRG**

(**A**) Gene set enrichment analysis (GSEA) across models (~ 0 + Batch + Diabetic.Type + X + Y + …) comparing DPN versus Diabetic donors for the Apoptosis hallmark gene set.

(**B**) Competitive gene set testing (CAMERA) for apoptosis, assessing enrichment of apoptosis-related genes between Diabetic and DPN groups.

(**C**) Apoptosis enrichment scores comparing Nageotte nodule burden for Diabetic-Low versus Diabetic-Mod-High groups.

(**D**) Heatmap of apoptosis-, stress-, and injury-linked proteins across donors, shown as row-scaled normalized abundance, with donor-level annotations indicating condition, pain/NeuP status, and Nageotte nodule burden (low/moderate/high).

**Supplementary Table 1**: Donor Demographics see attached excel file
